# Nucleosome composition regulates the histone H3 tail conformational ensemble and accessibility

**DOI:** 10.1101/2020.06.26.172072

**Authors:** Emma A. Morrison, Lokesh Baweja, Michael G. Poirier, Jeff Wereszczynski, Catherine A. Musselman

## Abstract

Sub-nucleosomal complexes including hexasomes and tetrasomes have been identified as intermediates in nucleosome assembly and disassembly. Their formation is promoted by certain histone chaperones and ATP-dependent remodelers, as well as through transcription by RNA polymerase II. In addition, hexasomes appear to be maintained in transcribed genes and could be an important regulatory factor. While nucleosome composition affects the structure and accessibility of the nucleosomal DNA, its influence on the histone tails is largely unknown. Previously, we found that the H3 tail accessibly is occluded in the context of the nucleosome due to interactions with DNA (Morrison et al, 2018). Here, we investigate the conformational dynamics of the H3 tail in the hexasome and tetrasome. Using a combination of NMR spectroscopy, MD simulations, and trypsin proteolysis, we find that the conformational ensemble of the H3 tail is regulated by nucleosome composition. Similar to what we previously found for the nucleosome, the H3 tails bind robustly to DNA within the hexasome and tetrasome, but upon loss of the H2A/H2B dimer, we determined that the adjacent H3 tail has an altered conformational ensemble, increase in dynamics, and increase in accessibility. Similar to observations of DNA dynamics, this is seen to be asymmetric in the hexasome. Our results indicate that nucleosome composition has the potential to regulate chromatin signaling at the histone tails and ultimately help shape the chromatin landscape.

## Introduction

The eukaryotic genome is packaged into the cell nucleus in the form of chromatin. The basic subunit of chromatin is the nucleosome, a complex of histone proteins and DNA. The canonical nucleosome core particle consists of ~147 base-pairs (bp) of DNA wrapped around an octamer containing one H3/H4 tetramer and two H2A/H2B dimers. In addition to this canonical species, sub-nucleosomal species, which contain fewer than eight histones, have been identified. These include the hexasome and tetrasome, which are lacking one or both H2A/H2B dimers, respectively (Figure 1A).

**Figure 1.**
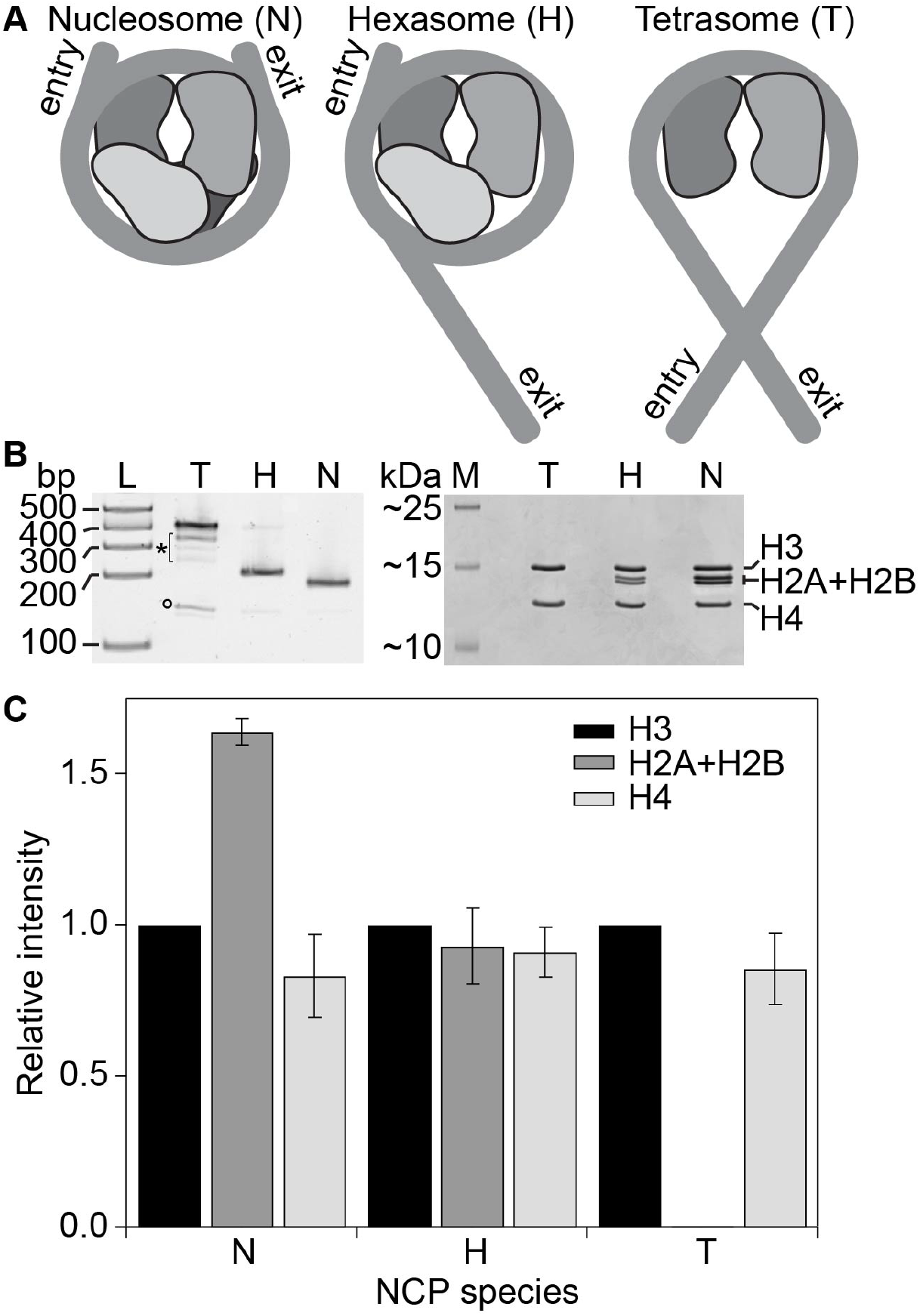
Composition of nucleosomal species. **A.** Cartoon depicting the composition of the three nucleosomal species investigated in this study, nucleosome, hexasome, and tetrasome. DNA and H3/H4 and H2A/H2B dimers are shown in shades of grey. Only the nucleosome core is represented. **B.** Gels characterize the purified nucleosome (N), hexasome (H), and tetrasome (T). Native PAGE (5% acrylamide, left) and denaturing SDS-PAGE (18% acrylamide, right) confirm the identity of the species. The native gel was visualized with ethidium bromide and includes TrackIt 100bp DNA ladder (L) for size reference. The denaturing gel was stained with Coomassie and includes Spectra BR marker (M) for size reference. The asterisk (*) denotes putative alternative positioning of the tetrasome, and the circle (°) marks free 601 DNA. **C.** The bar graph shows the relative intensities of the four histones within the three nucleosomal species as quantified from 18% denaturing acrylamide gels. The intensities (volumes) of gel bands were normalized to H3 to provide a relative intensity, and the average and standard deviation are depicted from four gel replicates.

For some time, these species have been studied *in vitro* and have been suggested to play a role in cellular processes such as transcription (reviewed in ^1^). Hexasomes and tetrasomes have been shown to be intermediates in chaperone-mediated and salt-dependent nucleosome assembly/disassembly^2–9^. In addition, hexasomes have been seen to form during transcription and ATP-dependent remodeling of nucleosomes^10–13^. Furthermore, the presence of hexasomes versus nucleosomes has been shown to differentially affect the activity of RNA polymerase II^14^ and the CHD1 chromatin remodeler^15,16^, supporting a regulatory role for sub-nucleosomes. Recently, these species have been observed *in vivo*^17,18^. It has been suggested that hexasomes exist as stable species near transcription start sites and may be an important regulatory factor.

A number of structural and biophysical studies have allowed for characterization of these species^6,19–26^. These studies have revealed that the histone core composition influences the DNA conformation and accessibility. Loss of an H2A/H2B dimer leads to unwrapping of ~30-40bp of DNA, which alters accessibility to digestion by endonuclease and transcription factor binding^15,19,22,27^. Notably, while the nucleosome and tetrasome are structurally pseudo-symmetric particles, the hexasome is structurally asymmetric both in the histone core and the associated DNA wrapping^1,6,15,20,22,23,27^. Intriguingly, DNA unwrapping and dimer loss have been observed to be asymmetric both *in vitro* and *in vivo. In vitro* studies reveal a dependence on DNA sequence, and *in vivo* this is correlated with transcriptional activity^27^. It has been proposed that this asymmetry may be important in reinforcing directional activity of RNA polymerase and chromatin remodelers^14,15,17,18^.

We recently proposed a structural model of the H3 tails in the context of the nucleosome. Based on nuclear magnetic resonance (NMR) spectroscopy and molecular dynamics (MD) simulations, we proposed that the H3 tails adopt a “fuzzy” complex with DNA^28–31^, interacting robustly but adopting a heterogenous and dynamic ensemble of DNA-bound states. This leads to occlusion of the tails and restricts access to histone tail binding domains^32^. This model suggests that chromatin signaling events could be regulated by modulating the DNA binding and conformational ensemble of the H3 tails. In the canonical nucleosome, the H3 tails protrude from between the two gyres of DNA near the entry/exit sites. Our previous MD simulations indicate that the tails form interactions with both gyres^32^. Thus, the loss of one or both H2A/H2B dimers and subsequent DNA unwrapping is predicted to significantly alter the conformational ensemble and possibly accessibility of the H3 tails.

Here, using a combination of NMR spectroscopy, MD simulations, and proteolysis assays we show that the H3 tails adopt unique conformational ensembles between nucleosome, hexasome, and tetrasome. Our results indicate that loss of H2A/H2B leads to an increase in the conformational dynamics of the H3 tail and accessibility to binding. Similar to the DNA dynamics, in the hexasome these effects are seen to be asymmetric. Together, these data suggest that conversion between nucleosome, hexasome, and tetrasome may modulate chromatin signaling at the histone tails and that this could function synergistically with concomitant changes in DNA accessibility.

## Results

### The H3 tail conformation is sensitive to nucleosome composition

To compare H3 tail conformational states between the canonical nucleosome core particle and the sub-nucleosome species hexasome and tetrasome, we used NMR spectroscopy. Nucleosome, hexasome, and tetrasome were reconstituted using H3/H4 tetramer containing ^15^N-labeled H3, and varied amounts of H2A/H2B dimer as required to obtain a given species (Figure 1, Figure 1—Figure Supplement 1). All species were reconstituted with the 147bp Widom 601 DNA (see methods section for details). Initial comparison of the ^1^H,^15^N-heteronuclear single quantum coherence (HSQC) spectra of each species (Figure 2A, Figure 2—Figure Supplement 1) reveals unique spectral attributes for the H3 tails within each species, indicating that nucleosome composition alters the conformation of the H3 tails.

**Figure 2.**
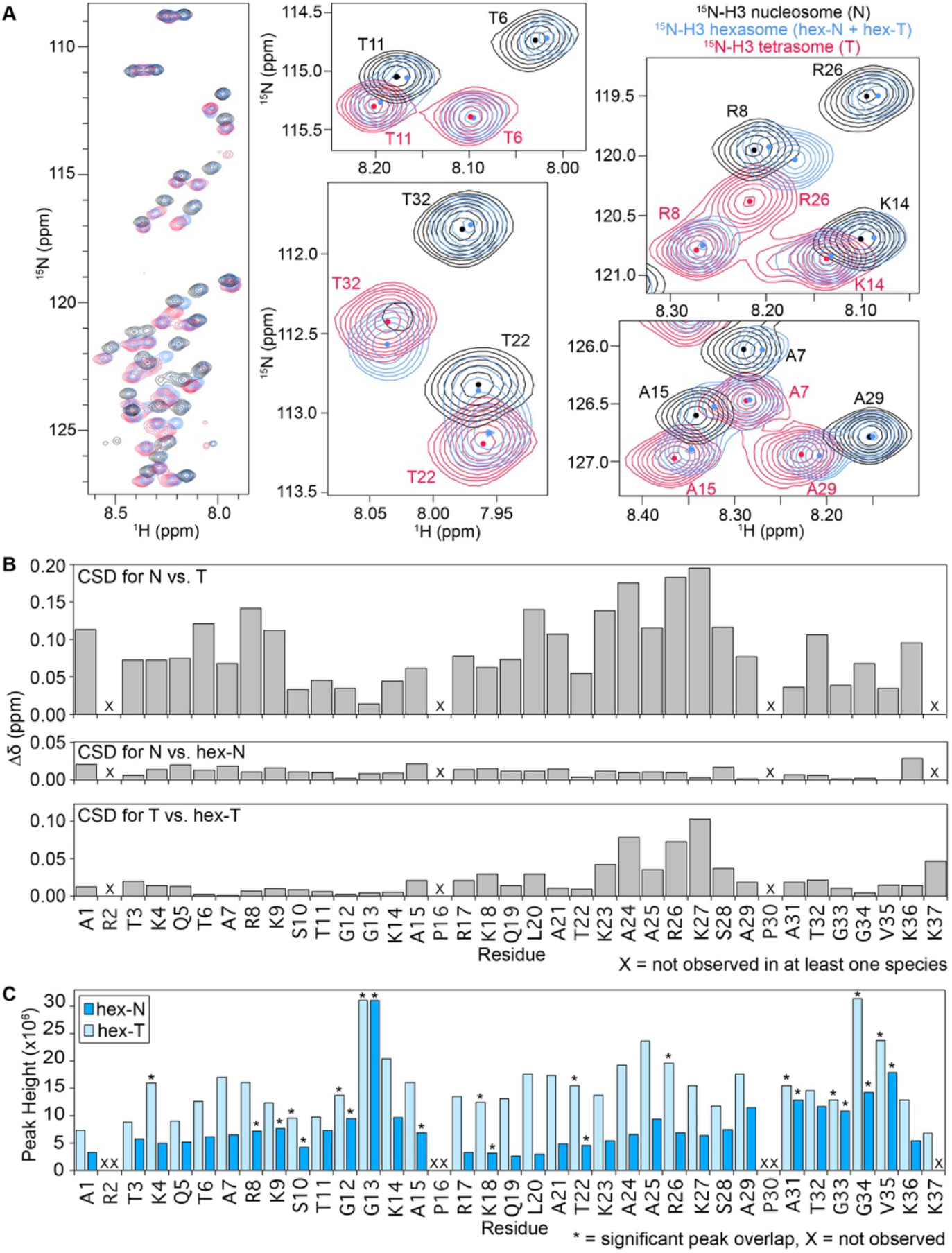
H3 tail conformation is distinct between nucleosomal species. **A.** Overlay of ^1^H/^15^N-HSQC spectra collected on ^15^N-H3-labeled versions of the three nucleosomal species, nucleosome (black), hexasome (blue), and tetrasome (red). Comparison of the spectra indicates that the H3 tail exists in different conformational ensembles between the nucleosome and tetrasome and suggests that hexasome contains one copy of H3 in a similar conformational ensemble as nucleosome and one copy of H3 in a similar conformational ensemble as tetrasome. Expanded regions of the overlay are shown for selected residues for closer comparison of histone tail states. Small circles mark the approximate center of each peak to aid in the spectral comparison. These spectra were collected on 44μM ^15^N-H3 NCP species in 20mM MOPS pH 7, 1mM EDTA, 1mM DTT, 7% D2O at 37°C and on an 800MHz spectrometer. **B.** Chemical shift differences (Δδ) between the nucleosome and tetrasome H3 tails (top), the nucleosome and hex-N H3 tails (center), and the tetrasome and hex-T H3 tails (bottom). This plot is shown as a function of H3 tail residue. **C.** Peak intensity (height) is plotted as a function of residue for the hex-N (blue) and hex-T (light blue) H3 tails within the hexasome. Residues that are not observed in the spectra are marked with an ‘X’. Residues with significant overlap that prevents accurate determination of peak height are marked by ‘*’.

One major difference between the spectra is in the number of peaks observed: 33 peaks for ^15^N-H3 nucleosome, 34 peaks for ^15^N-H3 tetrasome and 65 peaks for ^15^N-H3 hexasome (Figure 2—Figure Supplement 1). To better understand these differences, we carried out backbone assignments of the resonances. We previously assigned the ^15^N-H3 nucleosome peaks to H3 tail residues 1-36, with only a single set of peaks observed for the two tails^32^. For ^15^N-H3 tetrasome, a single set of peaks was also observed for residues 1-36, and an additional peak was observed corresponding to Lys37 (Figure 2—Figure Supplement 1). The single set of peaks for both ^15^N-H3 nucleosome and ^15^N-H3 tetrasome indicates that the two H3 tails within each of these nucleosomal species experience largely the same chemical environment (Figure 2A, black and red spectra), which is consistent with the structural pseudo-symmetry within each of these species.

Assignments for ^15^N-H3 hexasome show that the 65 observed peaks all correspond to the H3 tails, but in contrast to the nucleosome and tetrasome, two peaks are observed for most residues in the H3 tail (Figure 2A, Figure 2—Figure Supplement 1, blue spectrum). This indicates two distinct states (or ensembles of states) of the tails within the hexasome. The multiple peaks could be explained by i) the two H3 tails experiencing distinct chemical environments or ii) interconversion of both of the H3 tails between two states that is slow on the NMR timescale. Importantly, Levendosky et al. elegantly showed that hexasomes reconstituted using the Widom 601 sequence preferentially assemble with the single H2A/H2B dimer at the TA-rich side of the DNA^15^. In addition, it has been shown that the 601 DNA asymmetrically unwraps from the histone core, becoming more accessible on the side of the particle lacking the H2A/H2B dimer^15,27,33^. Thus, we hypothesize that the two sets of peaks arise from each of the H3 tails adopting a unique conformational ensemble, dependent on the presence or absence of the adjacent H2A/H2B dimer.

Additional insight into the conformations of the H3 tails within the different nucleosomal species can be gained by comparing chemical shifts of H3 tail resonances between nucleosome, hexasome, and tetrasome (Figure 2B and Figure 2—Figure Supplement 2). Overlay of spectra for the nucleosome and tetrasome reveals that, even though the number of peaks is the same, there are substantial differences in the chemical shift of all residues (Fig. 2A, compare black and red spectra). This reveals that loss of both H2A/H2B dimers leads to a change in the chemical environment of the H3 tails. Overlay of the hexasome spectrum reveals something quite striking: in the spectrum for ^15^N-H3 hexasome, half of the peaks overlay well with the nucleosome spectrum and the other half overlay well with the tetrasome spectrum (Figure 2A). Furthermore, these two sets of peaks correspond to residues of a full H3 tail (i.e. correspond to residues H3 1-36 or 37). Together, this leads us to hypothesize that one H3 tail in the hexasome adopts a conformation similar to the nucleosome and the other adopts a conformation similar to the tetrasome. As such, these will be referred to as the hex-N and hex-T tails, respectively.

To better quantitate these comparisons, chemical shift differences (CSDs or Δδs) between the nucleosome and tetrasome peaks, the hex-N and nucleosome peaks, and the hex-T and tetrasome peaks were calculated (Figure 2B). Peaks for the nucleosome versus tetrasome had an average Δδ=0.09. The majority of resonances had Δδ>0.05, with the largest differences observed for residues K23-S28, indicating substantial differences in conformation. In contrast, peaks for the hex-N tail as compared to the nucleosome have Δδ<0.03 along the entire length of the tail, indicating a highly similar conformation. While compared to the tetrasome, the majority of the hex-T tail peaks also have Δδ<0.03, there are some residues that exhibit greater differences from the tetrasome tail. In particular, residues K23-S28 and K37 have Δδ>0.03. These differences may reflect known differences in overall stability, positioning, and dynamics of the tetrasome^1,6,22,33^. In addition, the chemical shift differences between the hex-N and tetrasome peaks, and the hex-T and nucleosome peaks show similar differences to those observed between nucleosome and tetrasome (average Δδ=0.09 and 0.07, respectively, Figure 2—Figure Supplement 2). This further supports the conclusion that one H3 tail adopts a nucleosomal-like state and the other adopts a tetrasomal-like state.

Altogether, these results strongly suggest that while the H3 tails adopt distinct conformational ensembles between nucleosome and tetrasome, the two tails are symmetric in both. In contrast, hexasome H3 tails are conformationally asymmetric, with one H3 tail adopting a nucleosome-like ensemble (hex-N) and one H3 tail adopting a tetrasome-like (hex-T) ensemble.

### Loss of the H2A/H2B dimer increases the dynamics of the H3 tail

Analysis of NMR spectra of each species also provides insight into the dynamics of the H3 tails. Notably, signal for an additional residue (K37) is observable in spectra of ^15^N-H3 tetrasome as compared to nucleosome. The appearance of Lys37 indicates that the tail is more dynamic near the particle core in the tetrasome relative to the nucleosome. Consistent with one tail being in a tetrasomal-like state in the hexasome only a single peak is observed for Lys37 in the ^15^N-H3 hexasome (see Figure 2—Figure Supplement 2A). Additional insight into the dynamics of the tails can be garnered from comparison of peak intensity between species. Peak intensity reports on intrinsic dynamics but is also influenced by overall tumbling. Thus, we focused on the two sets of peaks in the hexasome for direct comparison because they are contained within the same particle. Therefore, the overall tumbling is internally controlled for since they have the same overall tumbling. Analysis of the intensities of the hexasome peaks reveals that the hex-T subset of peaks is on average 2.4-fold more intense than the hex-N subset of peaks. This difference is the largest (on average 2.6-fold) for peaks corresponding to the first 29 residues of the H3 tails (Figure 2C). This suggests that the hypothesized tetrasomal H3 tail is more conformationally dynamic than the hypothesized nucleosomal H3 tail.

In a previous study, we concluded that the H3 tails are robustly but dynamically associated with DNA in the context of the nucleosome. This was concluded in part because the NMR spectra of the nucleosomal state was shifted from that of a dynamically unrestricted H3 tail peptide in a manner consistent with binding to DNA^32^. To further investigate the conformational state of the hexasome tails, amide chemical shifts were compared between the hexasome and a peptide corresponding to the H3 tail (residues 1-44) (Figure 3). Overlay of the ^1^H,^15^N-HSQC/HMQC spectra for the hexasome and peptide reveals that, in general, the hex-T H3 peaks lie along a near-linear trajectory between the nucleosome (or hex-N) and peptide peaks, though not fully reaching the peptide chemical shifts. This suggests that, upon loss of the H2A/H2B dimer, the conformational equilibrium of the H3 tail is shifted towards a more conformationally unrestricted state (Figure 3). Notably, the hex-T chemical shifts are highly similar to chemical shifts for the H3 peptide bound *in-trans* to DNA or *in-trans* to a tailless NCP (Figure 4 of ^32^, Figure 3—Figure Supplement 1). Based on this, we hypothesize that the tetrasomal state of the H3 tail is still bound to DNA, but more conformationally dynamic than the nucleosomal state of the tail. We further hypothesize that this is due to unwrapping and subsequent lowering of DNA density near the tail, suggesting that the H3 tails sample a conformational ensemble that is linked to the conformation of the DNA.

**Figure 3.**
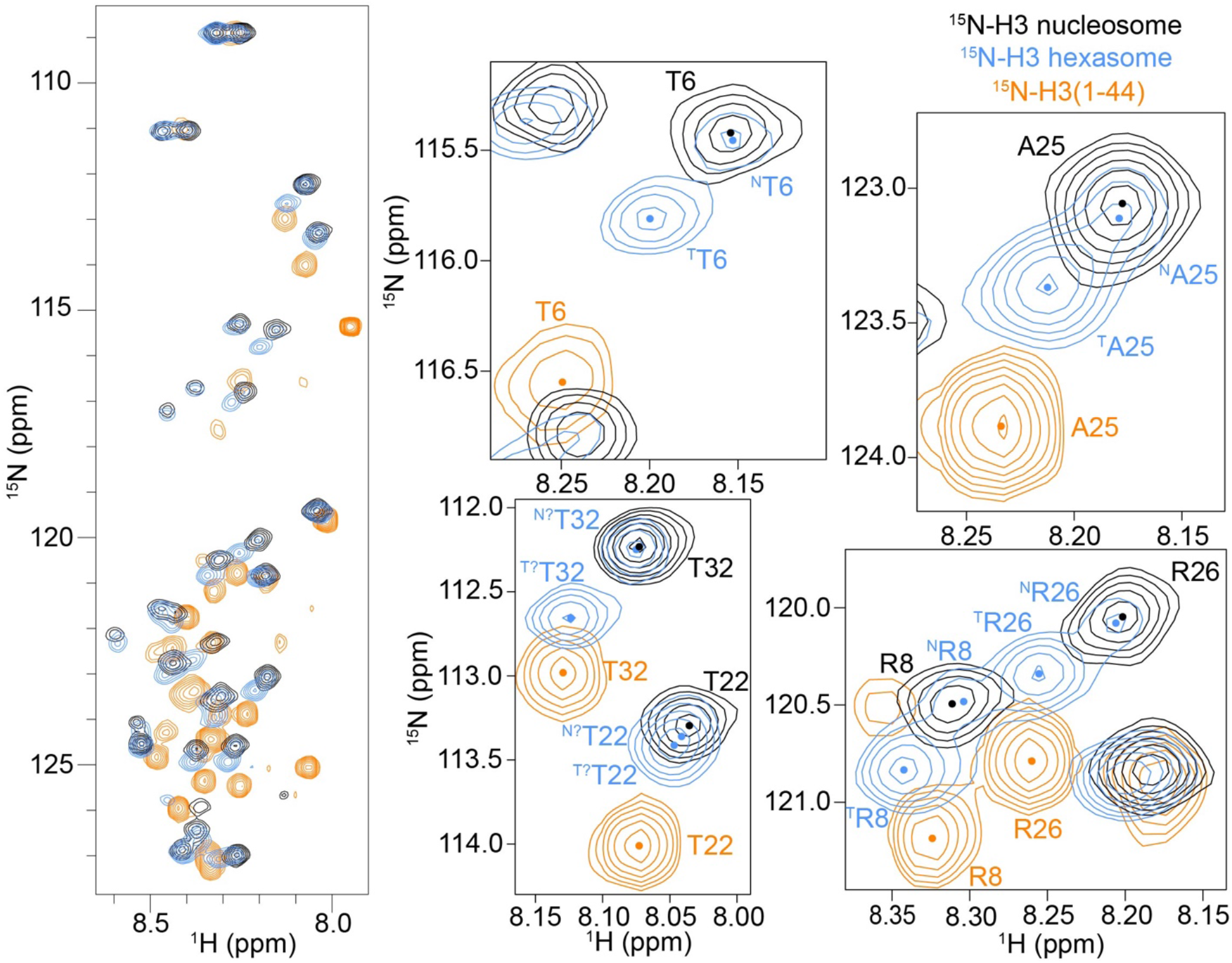
H3 tail conformation within tetrasome more closely mirrors H3 tail peptide. Overlay of ^1^H/^15^N-HSQC/HMQC spectra collected on ^15^N-H3 nucleosome (black), ^15^N-H3 hexasome (blue), and ^15^N-H3(1-44) (orange). Comparison of the spectra shows that, in general, the tetrasomal H3 tail experiences a more similar chemical environment to the H3 tail peptide than does the nucleosomal H3 tail, which suggests a more extended tail ensemble within the tetrasome than the nucleosome. Expanded regions of the overlay are shown for selected residues for closer comparison of histone tail states. Small circles mark the approximate center of each peak to aid in the spectral comparison. These spectra were collected on 44μM ^15^N-H3 nucleosome species or 110μM ^15^N-H3(1-44) in 20mM MOPS pH 7, 150mM KCl, 1mM EDTA, 1mM DTT, 7% D2O at 25°C and on an 800MHz spectrometer.

**Figure 4:**
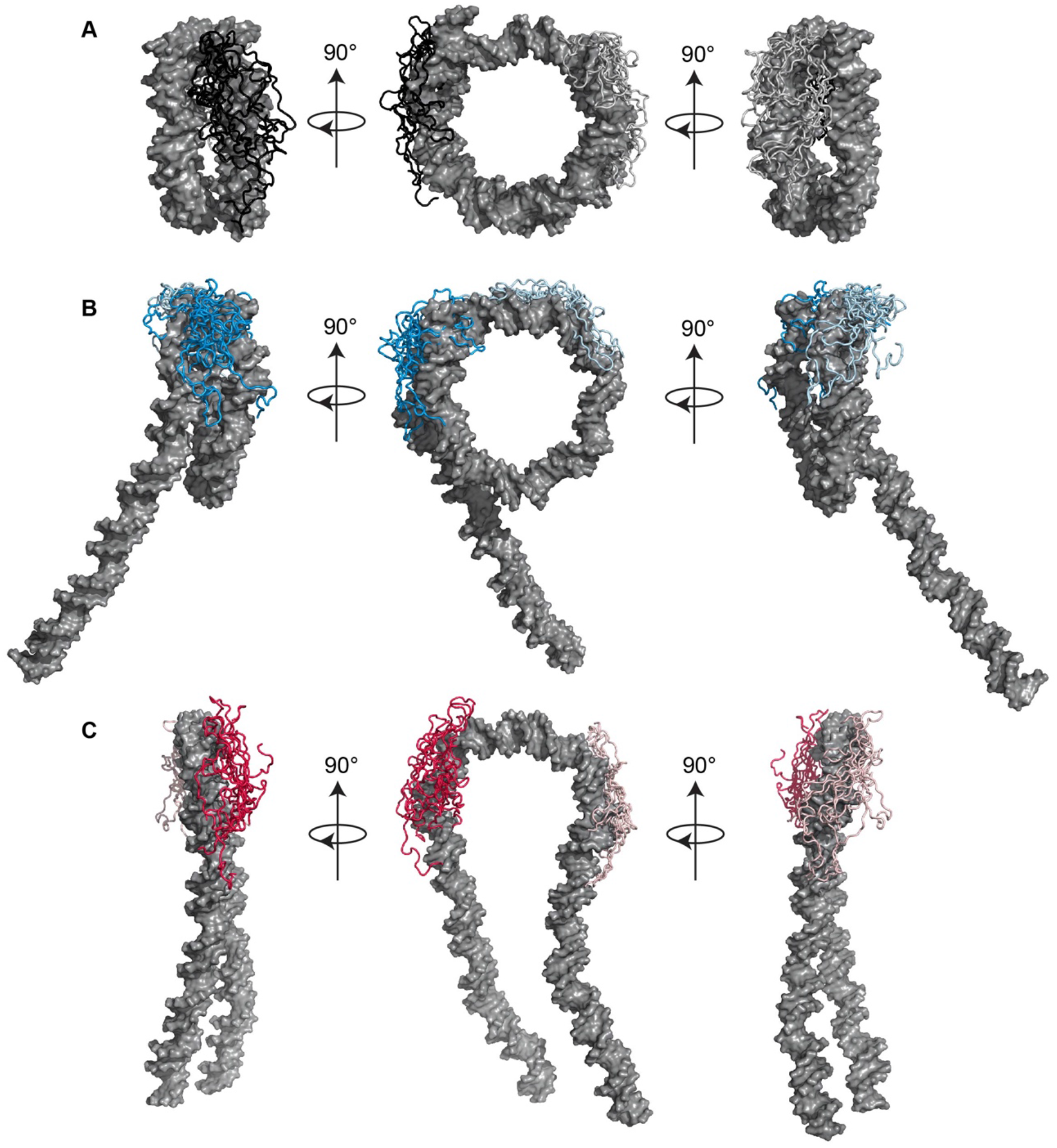
End states of nucleosomes and subnucleosomes obtained from simulations. End states of the H3 tails from ten simulations are shown on DNA from a single simulation for each of (A) nucleosome, (B) hexasome, and (C) tetrasome. The core histones and other histone tails are present in the simulations but removed in the figure for ease of visualization. Tail1 or hex-N is in the darker shade of each color (black, blue, and red) while tail2 or hex-T is in the lighter shade.

### Loss of dimer increases H3 tail conformational fluctuations in MD simulations

To further investigate the conformation and dynamics of the H3 tails in sub-nucleosomes, 10 × 250 nanosecond (ns) all-atom molecular dynamics (MD) simulations were carried out on nucleosome, hexasome, and tetrasome. We previously observed that the H3 tails in the nucleosome quickly adopt a DNA-bound state no matter their starting conformation. However, multiple DNA bound states were observed across several simulations with little energetic difference between them. Combined with NMR data, this led us to propose that the H3 tails adopt a fuzzy complex with DNA in the nucleosome context, interacting robustly but adopting a heterogenous and dynamic ensemble of DNA-bound states. In agreement with NMR data, we observe that in all simulations of the hexasome and tetrasome, the H3 tails bind to the DNA within 100 ns. Analysis of the end-state of all simulations reveals that in the nucleosome, hexasome, and tetrasome, the H3 tails adopt a heterogeneous ensemble of DNA-bound states (Figure 4, Figure 4—Figure supplement 1).

To assess the conformational dynamics of these DNA-bound states, the average root mean square fluctuation (RMSF) of Cα atoms of each tail over the 10 simulations for each species was calculated. These report on the dynamics of the tails with respect to the core. For all tails, average RMSF values were substantially greater than RMSF values for residues in the histone core indicating greater relative conformational dynamics (Figure 5). In the nucleosome, the two H3 tail RMSFs are similar with mean values of 3.0-6.0 Å, indicating a similar degree of conformational dynamics of each tail. In contrast, the initial portion of the H3 core (the α_1_ helix, residues 44-55) has average fluctuations of 0.7 Å. In the tetrasome, both H3 tails also have a similar degree of dynamics, but are substantially more dynamic than the nucleosome tails with calculated RMSFs between 5.5-10.0 Å. This indicates that removal of the H2A/H2B dimer increases the conformational dynamics of the H3 tails. Fluctuations in the H3 α_1_ helix were also increased to, on average, 1.8 Å. The calculated RMSF values for the hexasome revealed that, in contrast to the tetrasome and nucleosome, there is a difference between the two tails. The RMSF values (3.0-4.5 Å) for the hex-N H3 tail are similar to values for the nucleosomal H3 tail, with the exception of residues A29-V35 at the end of the tail. In contrast, the hex-T H3 tail shows a significant increase in flexibility (mean values of 3.5-5.5 Å). Similar to the tetrasome, the hex-T H3 core region also has a marked increase in flexibility, with RMSF values of 3.0 Å. Altogether this indicates that removal of the H2A/H2B dimer leads to an increase in the H3 core and tail dynamics, and that in the hexasome this introduces asymmetry.

**Figure 5:**
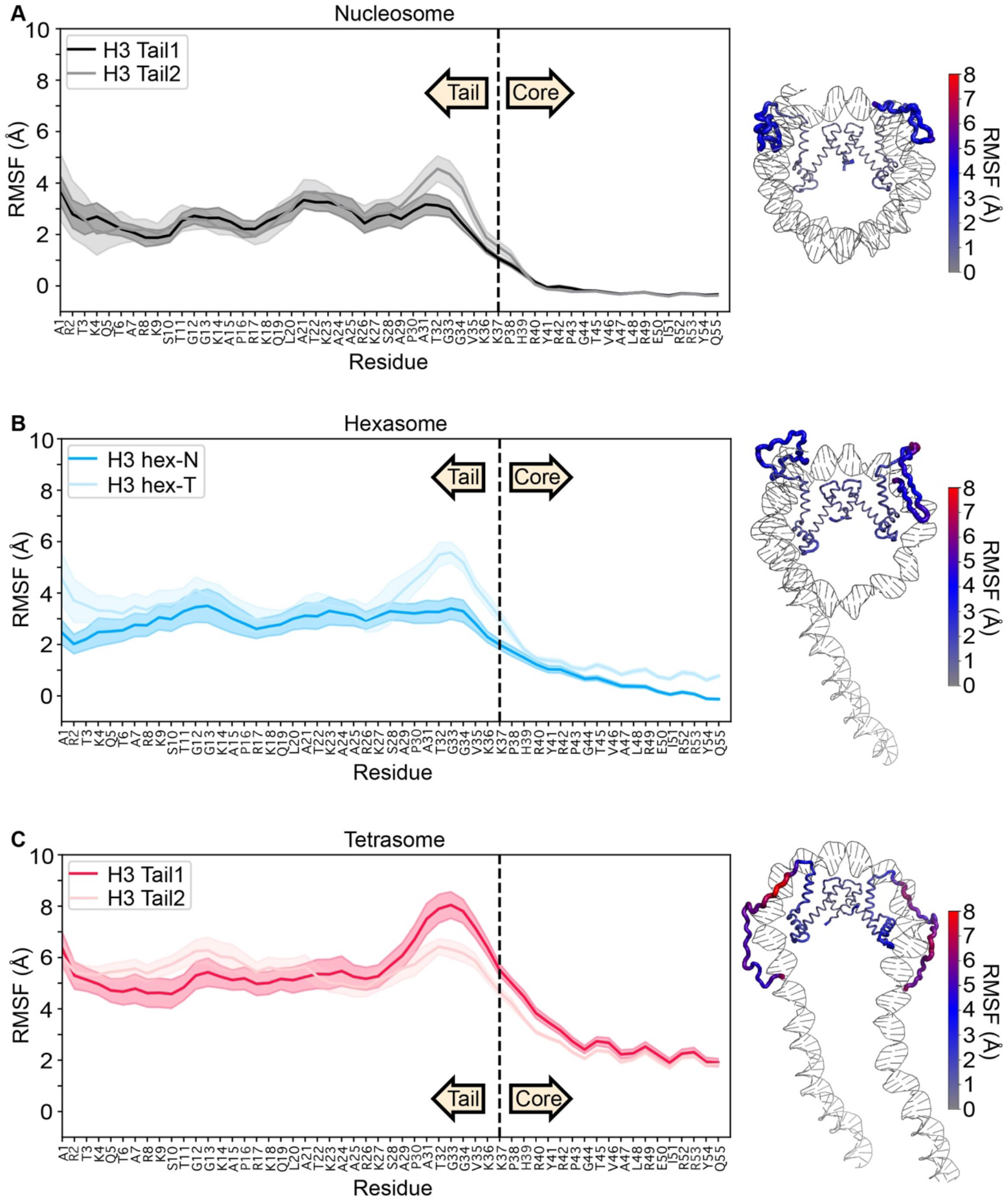
Residue-wise root mean square fluctuation (RMSF) values obtained from the equilibrated portion of MD trajectories. Plots are for (A) nucleosome, (B) hexasome, and (C) tetrasome with the average of ten simulations plotted as a solid line and the standard error of the mean shaded. Data are plotted for the first 55 residues of H3, where residues 1-37 and 38-55 are defined as tail and the initial region of the core, respectively. RMSF values are also plotted on a representative end-state structure of each nucleosomal species. The thickness and color (see key) of the cartoon backbone represents the RMSF value for a given residue, where thicker indicates larger RMSF. H3 is the only histone displayed in the image for ease of visualization.

To further ascertain the nature of these increased dynamics, the individual dihedral motions across all 10 simulations for each species were quantified with Kullback-Leibler divergence calculations^34^. Results show minimal statistically-significant differences between the nucleosome, hexasome, and tetrasome tails (result not shown). Therefore, while RMSF calculations show that global motions with respect to the core are increased in the hex-T and tetrasomal H3 tails, the differences in localized dihedral motion appear to be relatively minor on the hundreds of nanoseconds timescale. Together, this reveals that increased fluctuations observed upon loss of the H2A/H2B dimer are largely due to increased sampling of conformational space relative to the core (which would include increased dynamics of the bound DNA itself^22^) but does not increase localized dynamics of the DNA-bound states.

### Dimer loss leads to more extended and solvent-exposed states of the H3 tail

To further understand the impact of dimer loss on the H3 tail conformational ensemble, the average inter-residue distances along the H3 tails were calculated, which report on the tail compactness (Figure 4—Figure supplement 2). Results show that in all systems, the H3 tails are devoid of any secondary structure elements, consistent with our previous NMR results and simulation studies^32^. Compared to the nucleosome, there are decreased long-range intramolecular contacts in the tetrasomal tails, indicating that the H3 tails adopt less-compact conformations (that is, they are more extended) upon loss of the H2A/H2B dimer (Figure 4—Figure supplement 2). In the hexasome, the two tails are conformationally asymmetric, with the hex-T tails resembling the tetrasome with fewer long-range intramolecular contacts as compared to the hex-N tails. Interestingly, the hex-N tails adopt even more compact conformations than the nucleosomal tails. Together, this analysis suggests that dimer loss and DNA opening modulate the conformational ensemble of the adjacent H3 tail towards more extended states along the DNA. While in the hexasome, the H3 tail of the wrapped side becomes more compacted.

To further quantify the conformational states of the H3 tails, contacts between the tails and DNA super helical locations (SHL) were calculated (Figure 6). For the nucleosome, the tails are seen to bind on either side of the dyad (SHLs −2.5 to 2.0), and outer DNA turns SHL −7.0 to SHL −5.0, and SHL 6.0 to SHL 7.0. Notably, this positioning is in agreement with a cross-linking study that found contacts between the H3 tail (probe placed at H3T6C or H3A15C) and SHLs ±1.5 and ±2.0 in nucleosomes formed with 207bp-601 DNA^35^. (Interestingly, this folding back of the tail to interact with a range of locations on the core DNA was observed even though linker DNA was present.) In comparison, the tetrasomal H3 tails extend away from the dyad, binding at SHLs −3.5 to 3.0, without making any contacts with SHL 0.0. This could be due to the DNA unwrapping, facilitating additional SHL contacts more distant from the dyad, and is consistent with a more extended conformation (Figure 4—Figure supplement 2). For the hexasome, the hex-N tail is similar to a nucleosomal conformation, forming contacts with inner and outer DNA turn around SHL −7.0 to −5.5, and SHL 0 to 2.0. In contrast, the hex-T tail adopts unique contacts with SHL −2.0 to 0.5, occupying the region on and near the dyad and even extending across the dyad to make some cross-gyre interactions (SHL −7.0) with the wrapped DNA. This is again consistent with a more extended conformation (Figure 4—Figure supplement 2). Notably, the loss of any contacts with SHL −7.0 to −5.5 is consistent with the previous observation that in hexasomes the unwrapped DNA is as accessible as naked DNA to transcription factor binding, indicating no competition with histone tails for binding to this DNA^27^. Of all species, the nucleosome tails have the highest number of contacts per base-pair, which are comparable to the number of contacts in the hex-N tail. In contrast, the hex-T tail forms the fewest number of contacts with the DNA.

**Figure 6:**
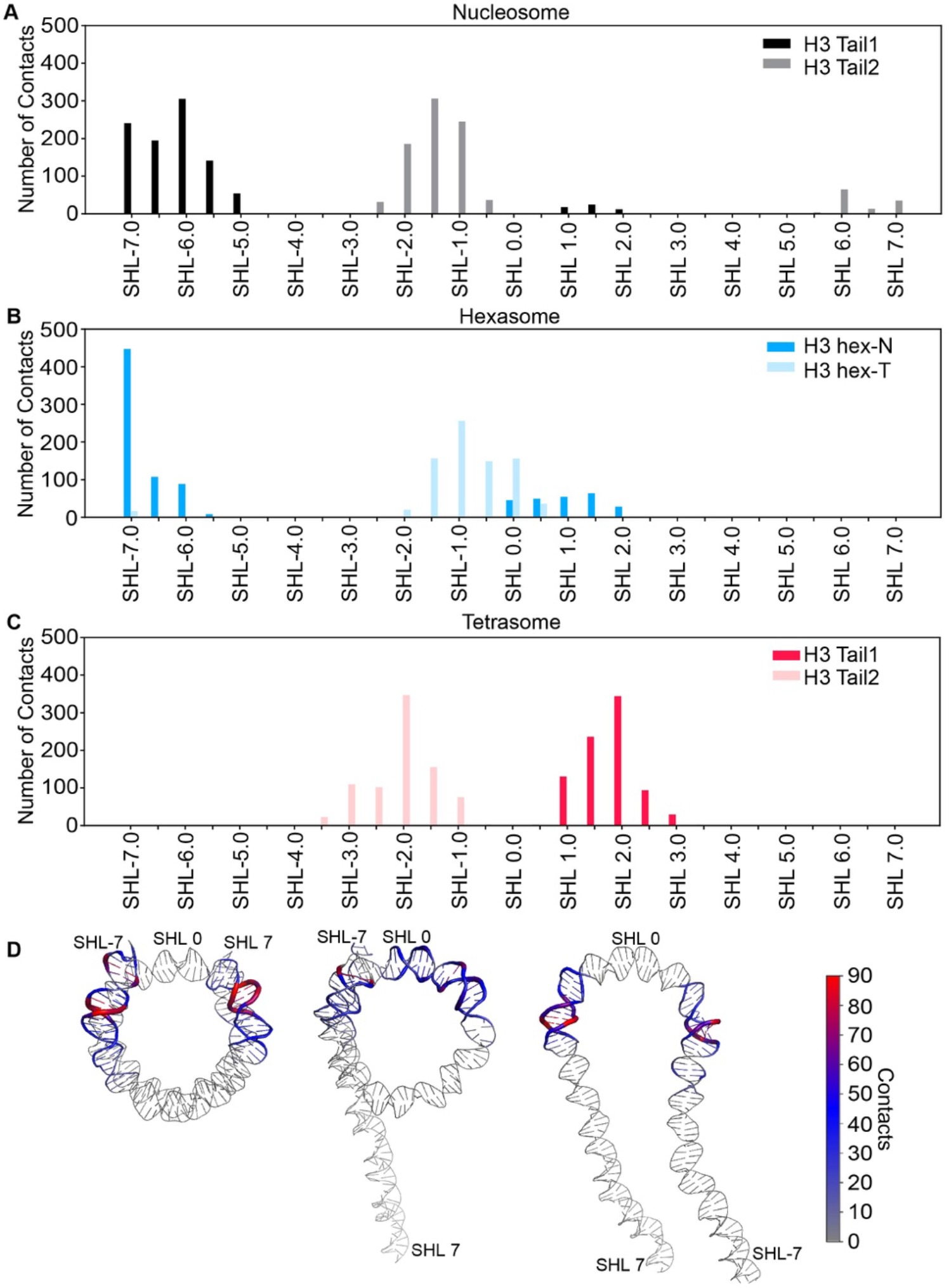
H3 tail contacts with DNA super helical locations (SHLs) in nucleosomal and subnucleosomal species. Plots are for (A) nucleosome, (B) hexasome, and (C) tetrasome. Contacts are summed over each SHL and plotted for each H3 tail. D. The number of contacts formed between H3 tail residues and DNA base-pairs were mapped onto nucleosomal and subnucleosomal DNA. The thickness and color (see key) of the cartoon backbone represents the number of contacts for a given base pair, where thicker indicates more contacts. A representative end-state structure is used for each species. Histones are omitted from the image for ease of visualization.

To analyze the interaction energetics of these H3 tail/DNA conformations, an MM/GBSA analysis was performed for residues 1-37 of the H3 tail. Nucleosomal H3 tails bound to DNA with similar energies of −131.0 ± 1.7 kcal/mol and −132.1 ± 1.6 kcal/mol, respectively (Table 1). In comparison the tetrasomal H3 tails bound to DNA weaker with energies of −119.0 ± 2.4 kcal/mol and −116 ± 1.3 kcal/mol, which are not statistically significantly different (p>0.2) from each other. In the hexasome, the hex-T tail bound with a statistically significantly (p<0.0001) lower energy than the hex-N tail at −110.5 ± 1.5 kcal/mol and −123.3 ± 2.0 kcal/mol, respectively. These weaker H3 tail binding affinities upon H2A/H2B dimer loss are accompanied by greater solvent exposure of these residues. This was determined from the calculated solvent accessible surface area (SASA), which is ~250 Å^2^ greater in the tetrasomal and hex-T tails as compared to the nucleosomal and hex-N tails. We note that care must be taken when interpreting MM/GBSA results, as it involves several approximations. These include a mean-field solvent and the lack of configurational entropy calculations, which is likely significantly different between the tails as demonstrated by RMSF calculations. Therefore, these energies should be interpreted only qualitatively.

**Table 1.**
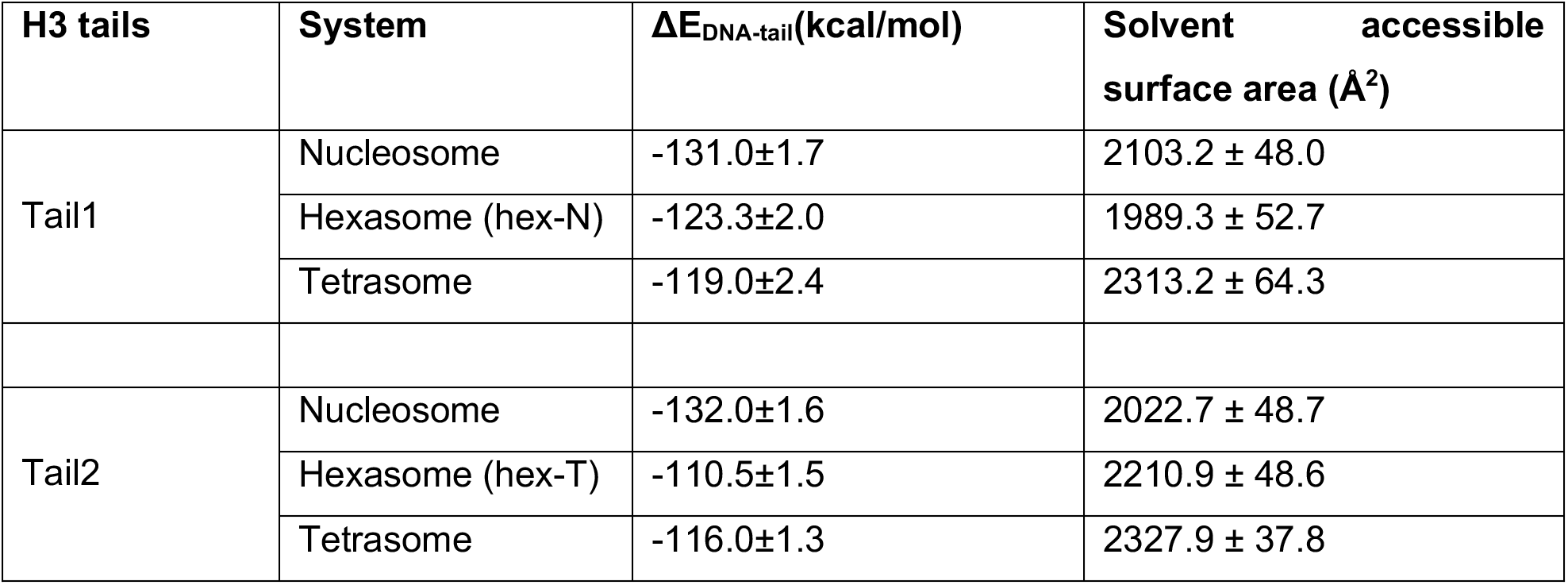
Binding energetics and solvent exposed surface area of individual tails from molecular dynamics simulations.

Altogether, the MD simulations are consistent with the NMR data, where upon loss of the H2A/H2B dimer and unwrapping of the DNA, the H3 tail remains in a DNA bound state. However, the tails experience increased conformational dynamics with respect to the core. In the hexasome, this leads to asymmetric conformational ensembles and dynamics of the two H3 tails.

### The H3 tail has differential accessibility between nucleosomal species

To test whether the increased dynamics in the H3 tail upon H2A/H2B dimer loss leads to increased accessibility, we performed trypsin proteolysis of the H3 tails in the context of the nucleosome, hexasome, and tetrasome. Trypsin proteolysis serves as a useful method to probe the general accessibility of the histone tails in a relatively non-sequence-specific manner because trypsin preferentially cleaves on the C-terminal side of lysine and arginine residues, which are spread out along the length of the tail^36,37^.

Each nucleosomal species was incubated for 20 minutes with three different amounts of trypsin (1:1/100, 1:1/500, and 1:1/2500 molar ratio of nucleosomal species:trypsin). The amount of proteolysis was ascertained via SDS-PAGE (Figure 7A, Figure 7—Figure Supplement 1A) by monitoring the amount of full-length H3 remaining. Note that since trypsin proteolysis can occur at multiple sites on the tails and we quantify the fraction of full length H3 remaining, this is a measure of the overall accessibility. In addition, the signal represents a sum of both H3s within each sample, since the two tails cannot be distinguished. Plotting the amount of full-length H3 remaining for each ratio of trypsin tested (Figure 7A) shows that each species undergoes different levels of proteolysis. The general trend indicates substantially greater proteolysis of the tetrasomal H3 tails as compared to the nucleosomal H3 tails. Notably, the level of proteolysis of the hexasomal H3 tails lies in between the nucleosome and tetrasome.

**Figure 7.**
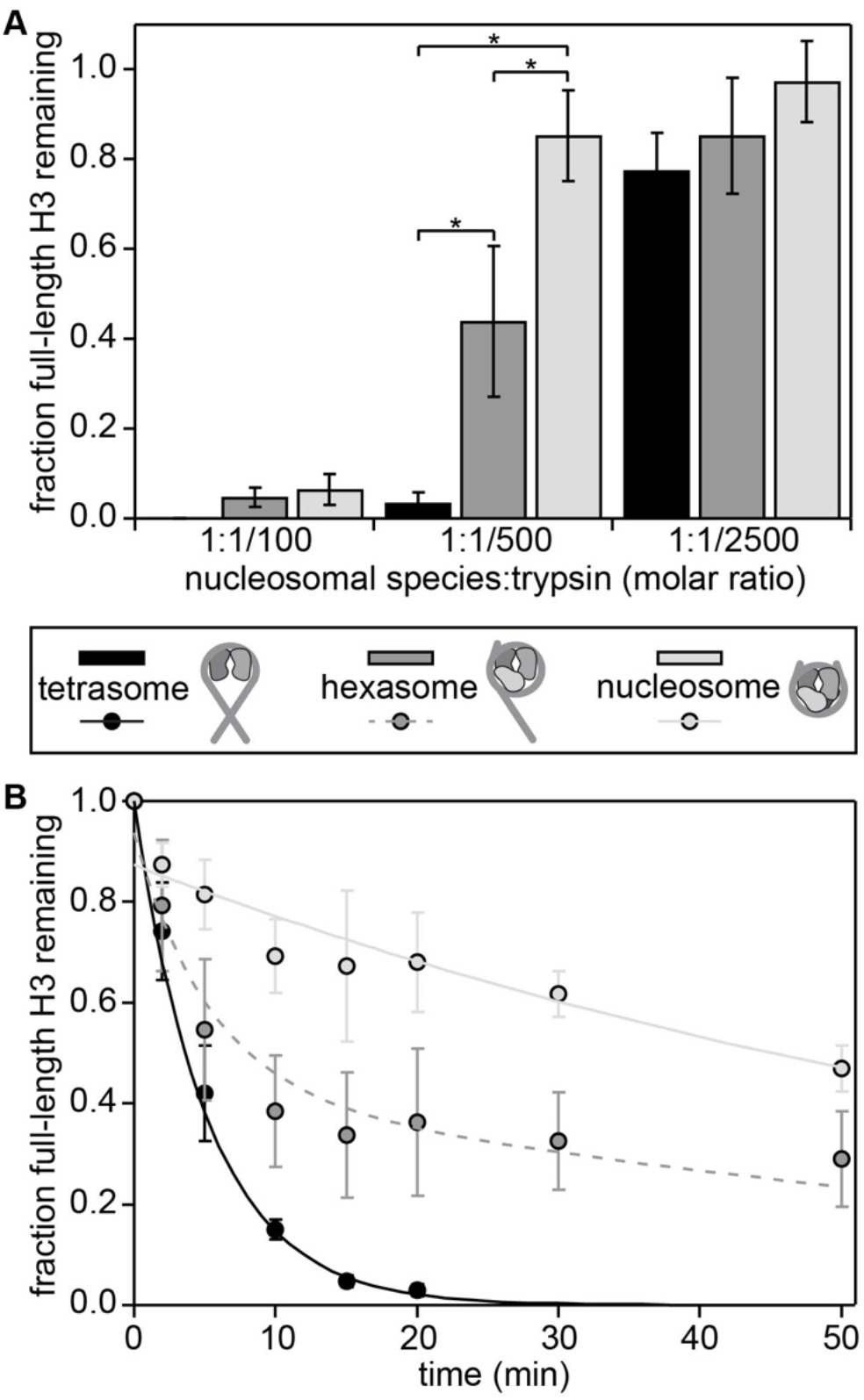
Trypsin digestion assays support differential accessibility of the H3 tail within different nucleosomal species. Gel-based trypsin digestion assays were used to probe tail accessibility. **A.** The bar graph displays the results from proteolysis at a constant concentration of nucleosomal species (3μM) and varying concentrations of trypsin. The progress of the trypsin proteolysis was assessed via SDS-PAGE. The fraction of full-length H3 remaining at t=20min as compared to t=0 is shown, as determined by the intensities (volumes) of gel bands for full-length H3. The average and standard deviation are depicted from three experimental replicates. Data are marked (*) if differences between nucleosomal species at a given trypsin concentration were determined to be statistically significant as determined by a two-way ANOVA followed by a tukey post-hoc analysis (p<0.05). **B.** The time course of a trypsin digestion was followed at the 1:1/500 molar ratio of nucleosomal species:trypsin with 3μM of the given nucleosomal species. The progress of trypsin proteolysis was assessed via SDS-PAGE. The fraction of full-length H3 remaining is plotted as a function of time. The intensities (volumes) of gel bands for full-length H3 were normalized to H3 at t=0, and the average and standard deviation are depicted from three gel replicates. Weighted single exponential fits (constrained to decay to zero and to have y-intercept ≤ 1) are shown for nucleosome and tetrasome (solid lines). The sum of the two exponential decays, with each weighted by one-half, represents the predicted time course for hexasome (dashed line).

To further quantify the H3 tail accessibility to trypsin digest, we acquired kinetic time courses at a ratio of nucleosomal species:trypsin of 1:1/500 (Figure 7B, Figure 7—Figure Supplement 1B). Experiments were conducted in triplicate, and the data were analyzed using a weighted fit of the fraction of remaining full-length H3 to a single exponential, similar to in ^38,39^ (see Materials and Methods for additional details). Importantly, native PAGE confirms that the nucleosomal species remained largely intact during the experiments (Figure 7—Figure Supplement 1). However, data was fit allowing for an initial offset on the y-axis to account for any small population of nucleosomes that may have fallen apart during the rapid mixing at the beginning of the digestion as is done in restriction enzyme digestions experiments^38^. The single-exponential fits imply cleavage rates of k_obs_=0.012±0.002 min^−1^ for nucleosome and k_obs_=0.19±0.02 min^−1^ for tetrasome (Figure 7B, solid light-grey and black lines, respectively). Under the conditions that the digestion rate is first order in enzyme (trypsin) concentration, which is support by Figure 7A and Figure 7—Figure Supplement 1A (lower left), the forward rate of digestion is proportional to the H3 tail site exposure equilibrium constant. This is the relative concentration of H3 tail accessible states as compared to inaccessible states. This overall approach is analogous to the studies that use restriction enzyme to measure DNA accessibility with partially unwrapped nucleosomes^40^. From the ratio of k_obs_, the relative site exposure probability is calculated as 15.8 ± 0.3, implying that the accessibility of the H3 tails is an order of magnitude greater upon loss of the H2A/H2B dimers. If the hexasome consists of one nucleosomal H3 tail and one tetrasomal H3 tail, as expected from the NMR data, the hexasome time course should be a sum of the two exponential decays, with each weighted by one-half (Figure 5B, dashed medium-grey line). Indeed, the experimental data for the hexasome are in very close agreement with this predicted time course (p=0.999 in a two-sample t-test), strongly supporting that one H3 tail is in the nucleosomal state and the other H3 tail is in the tetrasomal state.

Altogether, these studies indicate that loss of the H2A/H2B dimer leads to increased accessibility to H3 tail binding proteins. In addition, it supports results from NMR and MD analysis that the hexasome contains one tail in a nucleosomal state and one tail in a tetrasomal state.

## Discussion

In this study, we find that the conformational ensembles and accessibility of the histone H3 tails are modulated by nucleosome composition. Taken altogether, NMR, MD, and proteolysis analysis support a model wherein loss of the H2A/H2B dimer leads to a shift in the conformational ensemble, increased conformational dynamics, and increased accessibility of the adjacent H3 tail (Figure 8). Specifically, our data support a model in which the H3 tail adopts a broader conformational ensemble along the length of the DNA and increased transitions between states. This could be due to a change in the density of DNA around the tail as dimer loss leads to DNA unwrapping. These results are in agreement with recent fluorescence studies that observed an increase in H3 tail dynamics upon salt-induced loss of dimer to form hexasome^26^. In addition, they are reminiscent of recent studies which found that binding of HMGN1 and HMGN2 to nucleosomes shift the location of H3 tail-DNA contacts, which was proposed to be involved in modulating chromatin condensation^35^. Thus, modulating the conformational ensemble of the H3 tail within the nucleosome (or its sub-species) may be a general mechanism for regulating chromatin structure and accessibility.

**Figure 8.**
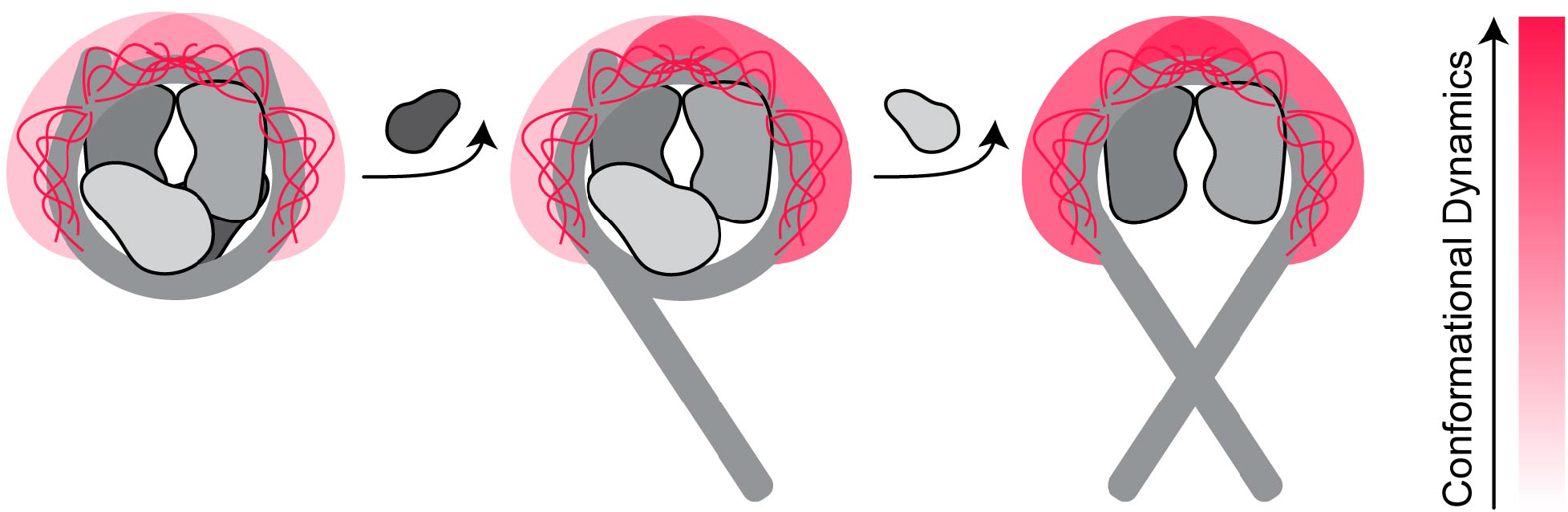
Model for the effect of nucleosome assembly state on H3 tail conformational ensemble. This cartoon model illustrates that loss of H2A/H2B dimer influences the conformational ensembles and dynamics of the H3 tails. DNA and H3/H4 and H2A/H2B dimers are shown in shades of grey. Of the histone tails, only the H3 tails are explicitly represented, and the cartoon depicts a cloud for the tail ensemble and explicitly depicts a subset of states within the ensemble. The shift in the color of the cloud from light to dark red indicates a shift toward faster dynamics within the conformational ensemble. The hexasome contains one nucleosomal H3 tail and one tetrasomal H3 tail, in accordance with the presence and absence, respectively, of the H2A/H2B dimer.

Notably, the modulation of H3 tail conformational dynamics and accessibility mirror previously observed changes in DNA dynamics and accessibility. Loss of the H2A/H2B dimer leads to DNA unwrapping and increased transcription factor association to an exposed consensus site to the level of free DNA^27^. In the hexasome, unwrapping of one side stabilizes the still-wrapped side, decreasing the unwrapping dynamics and association of transcription factors to a consensus site as compared to a nucleosome. It has been hypothesized that this reduced DNA unwrapping is due to rearrangement of the histones upon loss of the dimer. Here, simulations suggest that in the hexasome the H3 tail of the still-wrapped side adopts a more compact conformation on the wrapped DNA. In addition, the H3 tail from the unwrapped side crosses the dyad to the still-wrapped side to make additional contacts with the wrapped DNA. Thus, the H3 tail may be aiding in the observed stabilization.

Previous studies addressing histone tail accessibility in the context of the nucleosome have shown that the tails are significantly occluded within the nucleosome as compared to histone peptides or refolded histones^32,39,41,42^. In the case of the H3 tail, accessibility to chemical modification is reduced by a factor of ~250 at 50mM NaCl and ~10 at 150mM NaCl^41^. It has been proposed that accessibility to the H3 tail could be modulated by a number of factors including histone PTMs and DNA dynamics. In this study, we find that, in the absence of salt, accessibility of the nucleosomal H3 tail is increased by a factor of ~16 upon H2A/H2B dimer loss. These results align with investigations into the linked PHD fingers of CHD4, which bind the H3 tails in an 80bp-tetrasome with greater affinity than the 147bp-nucleosome^41^. This suggests that nucleosome composition will modulate accessibility of chromatin-associated proteins to the H3 tail.

In addition, since loss of H2A/H2B dimer leads to both DNA unwrapping and site exposure, as well as increased H3 tail accessibility, these concomitant changes could function cooperatively. For example, a protein domain that binds transiently to partially unwrapped nucleosomal DNA is anticipated to increase the H3 tail accessibility to an H3 tail binding domain. This could result in cooperative binding similar to how adjacent transcription factor binding sites can result in cooperative binding^43^. Furthermore, if the DNA and H3 tail binding domains are within the same protein or complex, the concomitant increase in accessibility of DNA and histone H3 tail could multiplicatively increase the binding probability. This could preferentially target complexes to the side of the hexasome that is missing the H2A/H2B dimer. Future studies will be needed to directly investigate these potential cooperativity mechanisms.

While recent data indicates the presence of sub-nucleosomes *in vivo*, their role is not yet fully understood. However, *in vitro* studies indicate that they modulate the activity of chromatin regulators (such as ATP-dependent remodelers) and RNA polymerase. In particular, the observed asymmetry of the hexasome has been hypothesized to play an important regulatory role. Here we observe that the histone H3 tail conformational dynamics and accessibility are regulated by the sub-nucleosome state and are asymmetric in the hexasome. We expect this will modulate the activity of chromatin modifiers and ATP-dependent remodelers, helping to shape the chromatin landscape, and may also contribute to regulation of transcription.

## Materials and methods

### Histone and DNA purification

Histones and 147bp Widom 601 DNA were expressed/amplified and purified as described in^32^.

### Mass spectrometry on histone samples

Electrospray ionization mass spectrometry was used to analyze the histones to confirm that there was no carbamylation as described in^32^.

### Generation of nucleosomes and subnucleosomes

Nucleosomes were largely reconstituted as described in^44^. Nucleosome reconstitutions were prepared with two variations—either (i) by refolding octamer with equimolar ratios of the histones H2A, H2B, H3 and H4 or (ii) by refolding tetramer (with equimolar ratios of H3 and H4) and dimer (with equimolar ratios of H2A and H2B) separately. Then, either (i) the octamer was mixed with 601 DNA at a 1:1 molar ratio or (ii) the tetramer, dimer, and 601 DNA were mixed together at a 1:2.2:1 molar ratio. Both mixtures were then desalted using a linear gradient from 2M to 150mM KCl over 36-48 hours, followed by dialysis against 0.5xTE. In our hands, refolding octamer together (via method (i)) results in a mixture of hexasome and nucleosome after the salt dialysis reconstitution while refolding tetramer and dimer separately (via method (ii)) results in finer control of the final sample. Samples were then purified with a 10-40% sucrose gradient, which separates residual free 601 DNA and hexasome formed from method (i).

Hexasome samples were made either by isolating hexasome from nucleosome reconstitutions carried out via method (i) or by following method (ii), except mixing tetramer, dimer, and 601 DNA at a molar ratio of 1:1.1:1. Similarly, tetrasome samples were made by following method (ii), except mixing tetramer and 601 DNA at a molar ratio of 1:1 in the absence of dimer. All reconstitutions were purified via sucrose gradient (BioComp Gradient Station, New Brunswick, Canada) (Figure 1—Figure Supplement 1). Although the DNA footprint of nucleosome, hexasome, and tetrasome are different, all three were prepared using the 147bp Widom 601 sequence in order to hold the total DNA content of the three species constant. It is also important to note that Levendosky et al. showed that hexasomes reconstituted using the Widom 601 sequence form a homogeneous population of oriented hexasomes, with the single H2A/H2B dimer preferentially assembling at the TA-rich side of the DNA^15^.

Native- and SDS-PAGE were used to assess the formation of nucleosome, hexasome, and tetrasome along with their histone compositions. Bands were visualized with ethidium bromide or Coomassie for native and denaturing gels, respectively. Gels were imaged using an ImageQuant LAS 4000 imager (GE Healthcare). With native-PAGE, the nucleosome runs as the most compact particle, followed closely by hexasome and then tetrasome (Figure 1B, left). This supports the model wherein first one and then both arms of DNA open up upon the loss of one or two dimers, respectively, and these changes would lead to more extended structures. Additionally, the nucleosome runs as the densest species on a sucrose gradient, again followed closely by hexasome and then tetrasome (Figure 1—Figure Supplement 1), which is again consistent with the structural models^1,22^. Notably, the tetrasome runs as a collection of bands on native-PAGE, with one major species. The basis of this is unknown, but could be due to differential positioning of the tetramer along the DNA and/or due to the presence of multiple tetramers on a single 147bp. Tetrasome was observed to be unstable in the presence of KCl, leading to the appearance of free 601 DNA via native-PAGE. Thus, tetrasome samples were only studied in buffers without salt added. SDS-PAGE confirmed the composition of the four histones within the final samples used for experiments (Figure 1B, right). The band density was used as a measure of intensity and was quantified using the ImageJ program (NIH). As in^15^, H2A and H2B were integrated together due to their lack of resolution. The intensities of gel bands were normalized to H3 to provide a relative intensity, and the average and standard deviation are taken from four gel replicates. Similar to that seen by Levendosky et al. ^15^, when the intensities of the gel bands are normalized to that of H3, the nucleosome contains nearly twice as much H2A and H2B as hexasome (Figure 1C).

Nucleosome concentrations were determined via UV-vis spectroscopy using the absorbance from the 601 DNA (calculated ε_260_=2,312,300.9 M^−1^cm^−1^). Samples were diluted into 2M KCl prior to concentration measurements in order to promote nucleosome disassembly for more accurate concentration determination.

### NMR spectroscopy data collection and analysis

To obtain backbone assignments for H3 within the context of subnucleosomes, HNCACB and CBCAcoNH spectra were collected on a 360μM ^13^C/^15^N-H3 hexasome sample (i.e. 720μM of H3) and a 130μM ^13^C/^15^N-H3 tetrasome sample (i.e. 260μM of H3) at 45°C and 37°C, respectively, using a Bruker Avance NEO 600MHz spectrometer. The HNCACB was collected with 32 scans and 88 and 68 total points in the ^13^C- and ^15^N-dimensions, respectively. The CBCAcoNH was collected with 24 (hexasome) or 32 (tetrasome) scans and 88 and 90 total points in the ^13^C- and ^15^N-dimensions, respectively. Assignments at 45°C on ^13^C/^15^N-H3 nucleosome were used from ^32^. Temperature titration was used to transfer assignments to 25°C and 37°C. Data were processed in NMRPipe ^45^ and assigned using CcpNMR Analysis ^46^. Assignments are summarized in Figure 2—Figure supplement 1 and Supplemental Table 1. Due to the repetitive and unstructured nature of the H3 tail, there is chemical shift degeneracy in some of the resonances. Associated assignment uncertainty is noted in Figure 2—Figure supplement 1 and Supplemental Table 1. Similar to ^32^, ^1^H/^15^N-HSQC spectra collected on H3K_C_4me3-hexasome were used to help confirm assignments of residues 3-9. Notably, the majority of peaks had degeneracy in C_α_ and C_β_ chemical shifts and thus could not be definitively assigned to one of the two copies of H3 within the hexasome. As noted in the results, these were categorized into subsets referred to as hex-N and hex-T according to amide chemical shift overlap with the nucleosome and tetrasome species, respectively. This is also noted in Figure 2—Figure supplement 1 and Supplemental Table 1.

^1^H-^15^N HSQC spectra were collected on ^15^N-H3 nucleosome, hexasome, and tetrasome samples. Samples were exchanged into 20mM MOPS pH 7, 1mM DTT, and 1mM EDTA (with some samples also containing 150mM KCl), and 7% D_2_O was added prior to data collection. The majority of data were collected on a Bruker Avance II 800MHz spectrometer with cryogenic probe. The spectra of 601 DNA-bound H3(1-44) were collected on a Bruker Avance Neo 800MHz spectrometer with cryogenic probe. To account for differences between instruments, referencing of an apo-spectrum of ^15^N-H3(1-44) was shifted until spectra overlaid between instruments, and referencing of the 601 DNA-bound spectrum was shifted by the same amount. All NMR data were processed in NMRPipe ^45^ and analyzed using CcpNMR Analysis ^46^. The chemical shift difference (Δ*δ*) between samples was calculated by:

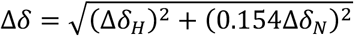

where *Δδ*_H_ and *Δδ*_N_ are the differences in the ^1^H and ^15^N chemical shift, respectively, between samples. Data plots were made in Igor Pro (Wavemetrics).

### Trypsin proteolysis assays

Trypsin proteolysis was used as a probe for site exposure on histone tails within nucleosomes and subnucleosomes. Digests were carried out on samples of reconstituted nucleosome, hexasome, and tetrasome at a fixed concentration of 3μM in 20mM MOPS pH 7, 1mM EDTA, and 1mM DTT.

Assays conducted at multiple ratios of trypsin were carried out at room temperature in 10μL reactions with 30nM, 6nM, and 1.2nM trypsin (Pierce product 90057, MS grade). Gel samples were taken prior to addition of trypsin (taken as t=0) and 20min after mixing with trypsin, when they were immediately mixed with 5x SDS loading dye and heated to 95°C for 10min. Gel samples contained 13pmol of the particular nucleosome species and were run on 18% tris-glycine SDS-PAGE gels followed by Coomassie staining. To check stability of the species over the course of the assay, 1.3pmol of the particular nucleosome species from before and after the assay were run on 5% native-PAGE gels and visualized with ethidium bromide.

Experiments with full time courses were conducted at 6nM trypsin (1:1/500 molar ratio of nucleosomal species:trypsin) in 80μL reactions. Samples were incubated in a thermomixer (Eppendorf) at 25°C while shaking at 350rpm. Gel samples were taken prior to addition of trypsin (taken as t=0) and at t=2, 5, 10, 15, 20, 30, and 50min after mixing with trypsin. Samples were quenched by immediately mixing with 5x SDS loading dye and heating to 95°C for 10min. Gel samples contained 15pmol of nucleosome or subnucleosome and were run on 18% tris-glycine SDS-PAGE gels and visualized with Coomassie stain. To check stability of the species over the course of the assay, 1.5pmol of nucleosome or subnucleosome from before and after the assay were run on 5% native-PAGE gels as before.

All experiments were run in triplicate. The native-PAGE confirmed that the nucleosomes, hexasomes, and tetrasomes remained largely intact over the course of the experiments.

### Analysis of trypsin proteolysis assays

Gel imaging—Gels were imaged using an ImageQuant LAS 4000 imager (GE Healthcare). The band density of full-length H3 was used as a measure of intensity and was quantified using the ImageJ program (NIH). The fraction of full-length H3 remaining at a given time was taken as the ratio of the band densities of full-length H3 at that time point and prior to the addition of trypsin.

Digests at multiple concentrations of trypsin—The amounts of full-length H3 remaining after 20min digestion at the three concentrations of trypsin were compared. To determine whether the extent of digestion was significantly different between the nucleosome and subnucleosomes at each concentration of trypsin, a two-way ANOVA followed by a tukey post-hoc analysis was run using R on the data sets that were collected in triplicate. A cutoff of p < 0.05 was used for significance.

Proteolysis kinetics—We treated the experimental data for site exposure on the H3 tails probed via trypsin proteolysis in the same manner as site exposure on DNA probed via restriction enzymes ^38,40^ and in a similar manner as site exposure on histone tails probed via chemical modification ^39,41^. These other experiments were designed for digestion and modification at single sites within the nucleosome. Although trypsin has many target sites within the histones, only the general proteolysis of the H3 tail is monitored by following the amount of full-length H3 remaining in the sample at a given time. The subsequent analysis makes several assumptions. First, we make the assumption that the system is in the limit of rapid conformational pre-equilibrium. In this limit, there is a first-order dependence of the observed rate constant (k_obs_) on enzyme concentration. Site exposure on nucleosomal DNA and H2B tails were shown to be in limit of rapid pre-equilibrium within the experimental contexts of ^40^ and ^39^, and the assumption of rapid pre-equilibrium was made for the H3 tail in ^41^. Thus, it is likely that site exposure on nucleosomal and subnucleosomal H3 tails is also in the limit of rapid pre-equilibrium in the proteolysis experiments described here. Although full kinetic data sets were only collected at a single concentration of trypsin, the single timepoint data collected at three trypsin concentrations suggests a linear relationship between k_obs_ (where the natural log of the fraction of full length H3 remaining is taken as a very rough proxy for k_obs_) and enzyme concentration (Figure 7—Figure Supplement 1A, lower left). An additional assumption is that the concentration of exposed histone tails is much less than the K_m_ of trypsin such that the free concentration of enzyme is equivalent to the total concentration of trypsin in the sample. Additional assumptions are that the concentrations of exposed H3 tail and H3 tail-trypsin complex are at steady state. Lastly, the assumption was made that the proteolysis events report predominantly on site exposure within the native conformation of the H3 tail within nucleosome or subnucleosome rather than site exposure that has been altered by a preceding cleavage event.

The average and standard deviation of the fraction of full-length H3 remaining at each time point was calculated from the triplicate data sets. The k_obs_ were determined from a weighted single exponential fit of the data average. The fit was additionally constrained to decay to zero and to have y-intercept ≤ 1. The y-intercept was allowed to be less than one to account for the possibility that the initial mixing of the sample led to dissociation of a subpopulation of particles. Under these constraints, the tetrasome experiment fit with a y-intercept of 1.0 ± 0.2 and the nucleosome experiment fit with a y-intercept of 0.87 ± 0.04. In studies of DNA site exposure, up to 10% of nucleosomes were observed to dissociate due to rapid mixing ^47^.

The ratio of site exposure equilibrium constants for tetrasome and nucleosome was taken as the ratio of the k_obs_ fit from the data sets for tetrasome and nucleosome. This only holds if the assumptions detailed above are valid. The error in the ratio of site exposure equilibrium constants was propagated from the error in the fits for the k_obs_ from the two data sets.

### Preparation/Generation of Canonical and Subnucleosomal particles for molecular dynamics simulations

Nucleosome models were constructed by taking a Widom 601 DNA molecule from PDB 3MVD sequence and mapping it onto the histone core coordinates from the 1KX5 PDB ^48,49^. Extended states of the H3 tails were built using MODELLER ^50^. Hexasome models were generated by removal of the H2A-H2B dimer from the nucleosome’s TA poor side, followed by implicit solvent molecular dynamics (MD) runs to create more open DNA structures ^22^. These initially involved position restraints on the first 107 bp of DNA and allowed the remaining 40 bp of DNA to relax for two ns in an implicit solvent environment with Watson-Crick base pair restraints. This geometry was then aligned with the histone hexamer to generate a crude hexasome intermediate. This was then simulated for 20 ns in an implicit environment to obtain a relaxed state with an extended DNA arm. Similarly, the tetrameric intermediate was generated by keeping only the (H3-H4)_2_ tetramer and allowing 40 bp of DNA from both sides of the nucleosome to relax during simulations. The starting tetrasome conformation had only ~66bp of DNA wrapped around the (H3-H4) tetramer, in accordance with the experimentally probed tetrasomal geometry ^22^. The initial conformations of the canonical and subnucleosomal particles are given in Figure 4—Figure supplement 1.

### Simulation Methods

All simulations were conducted in the CUDA-enable PMEMD engine of the AMBER software suite (v18)^51,52^. The Amber 14SB and BSC1 forcefields parameters were used for the protein and DNA respectively^53,54^. Implicit simulations were performed using mbondi3 and igb=8 ^55^. For explicit solvent simulations, all systems were neutralized and solvated with TIP3P waters and a 0.15 M KCL environment^56,57^. A 4-fs time-step was used in conjunction with SHAKE and hydrogen mass repartitioning for all simulations^58,59^. All systems were energy minimized for 5000 steps with a solute harmonic restraint of 10 kcal/mol/Å^2^, followed by 5000 steps with no restraints. For equilibration, we first performed 100 ps of constant volume simulations while the temperature was gradually heated from 10 K to 300 K. Then, the heavy atoms restrain were gradually released over 500 ps of NPT run. In explicit solvent simulations, pressure was controlled via a monte carlo barostat with a target pressure of 1 atm and a relaxation time of 3.0 ps^−1^. Production runs were performed at 300K using a Langevin thermostat^60^. We performed ten, 250 ns simulations per system in the NPT ensemble, accumulating 7.5μs of sampling across all the three systems. Trajectories were recorded every 10 ps and visualized using VMD^61^ and PyMol^62^. Analysis was performed on the last 150 ns of the simulations, allowing for 100 ns of equilibration.

### Simulation analyses

Root mean square fluctuation (RMSF) analysis was performed on the H3 proteins, the first 37 residues defined as the tails, and residues beyond this defined as the core region. Translations and rotations were removed by least squares fitting the backbone of the H3 and H4 histone, and RMSFs were computed on the Cα atoms. Reported RMSFs are the average of all 10 simulations. Errors are presented as the standard error of the mean obtained from 10 samples for each tail. Kullback-Leibler Divergence was performed using internal coordinates to compare conformational ensemble across the systems ^34^. Contacts analyses between H3 tails and DNA was performed using MDanalysis ^63^ and in-house python scripts, where contacts were defined as between heavy atoms of the H3 tail residues that were within a distance of 4.5 Å from DNA heavy atoms. Interaction energies between the H3 tails and DNA were determined from the sum of tail residue contributions to DNA binding via an MM-GBSA (Molecular Mechanics Generalized Born Surface Area) analysis with igb=5 and a salt concentration of 0.15M ^64^. Error bars represent the standard error of the mean, with a decorrelation time of 10 ns which is based on a statistical inefficiency test of MM/GBSA values.

---

**Supplementary Table 1.** Summary of assigned chemical shifts for nucleosome, hexasome, and tetrasome. See also Figure 2—Figure supplement. Due to the repetitive and unstructured nature of the H3 tail, there is chemical shift degeneracy in some of the resonances. Assignment uncertainty is noted in this table. For hexasome, the majority of peaks had degeneracy in C_α_ and C_β_ chemical shifts and thus could not be definitively assigned to one of the two copies of H3.

**Figure 1—Figure supplement 1.**
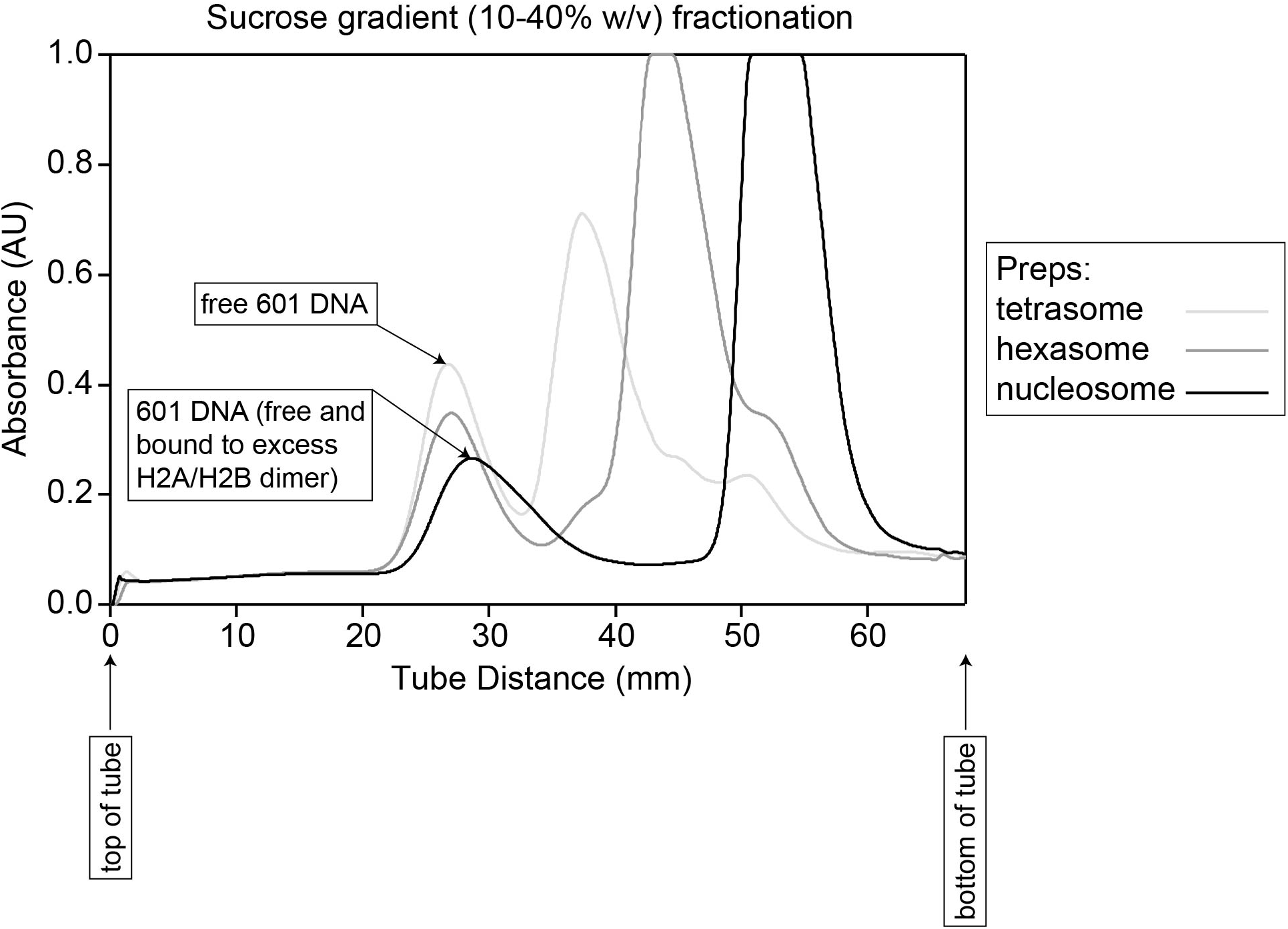
Sucrose gradient purification of nucleosomes and subnucleosomes. The reconstitutions of nucleosomes and subnucleosomes (described in materials and methods) were purified via sucrose gradient (10-40% w/v sucrose). The major peak for each was used for experiments.

**Figure 2—Figure supplement 1.**
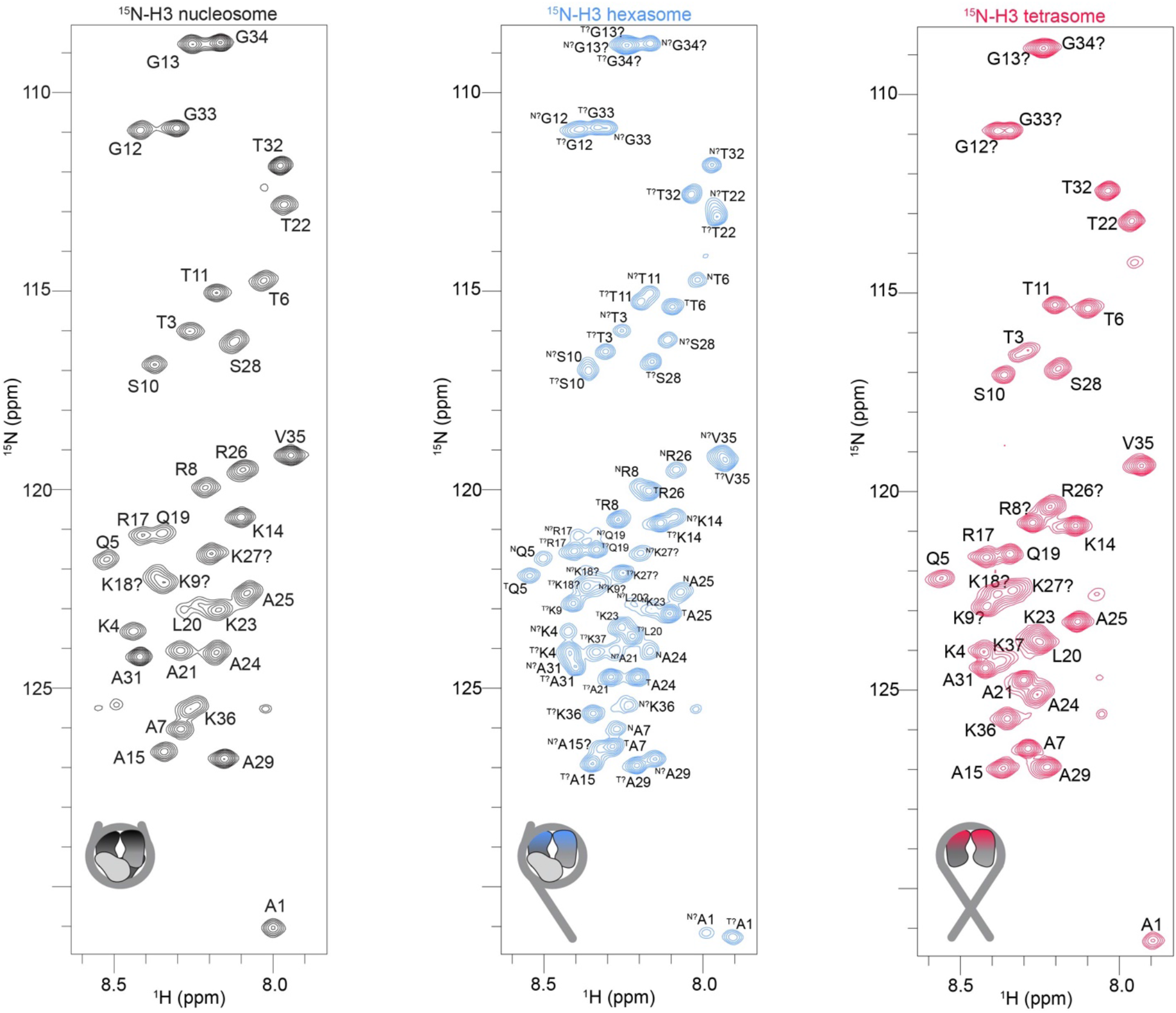
Full-sized ^1^H/^15^N-HSQC spectra of ^15^N-H3-labeled nucleosome (black), hexasome (blue), and tetrasome (red) used in main text Figure 2. Peaks are labeled with residue assignments, and assignment uncertainty is indicated by ‘?’ (see Supplemental Table X1). For hexasome, hex-N and hex-T tail assignments are indicated by the super-script N and T designations (along with uncertainty status).

**Figure 2—Figure supplement 2.**
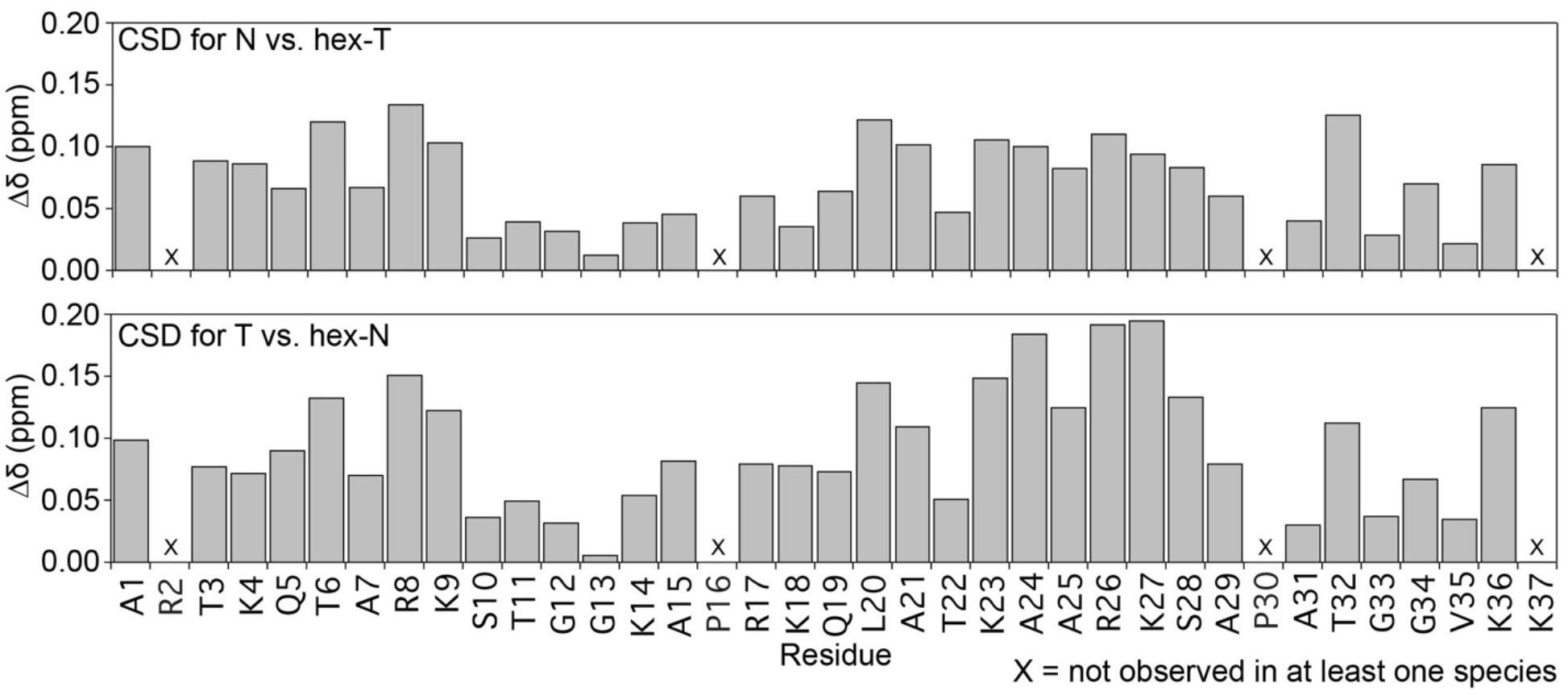
Chemical shift differences (Δδ) between the nucleosome and hex-T H3 tails (top) and the tetrasome and hex-N H3 tails (bottom). These plots are shown as a function of H3 tail residue. Residues that are not observed in the spectrum of at least one species are marked with an ‘X’.

**Figure 3—Figure supplement 1.**
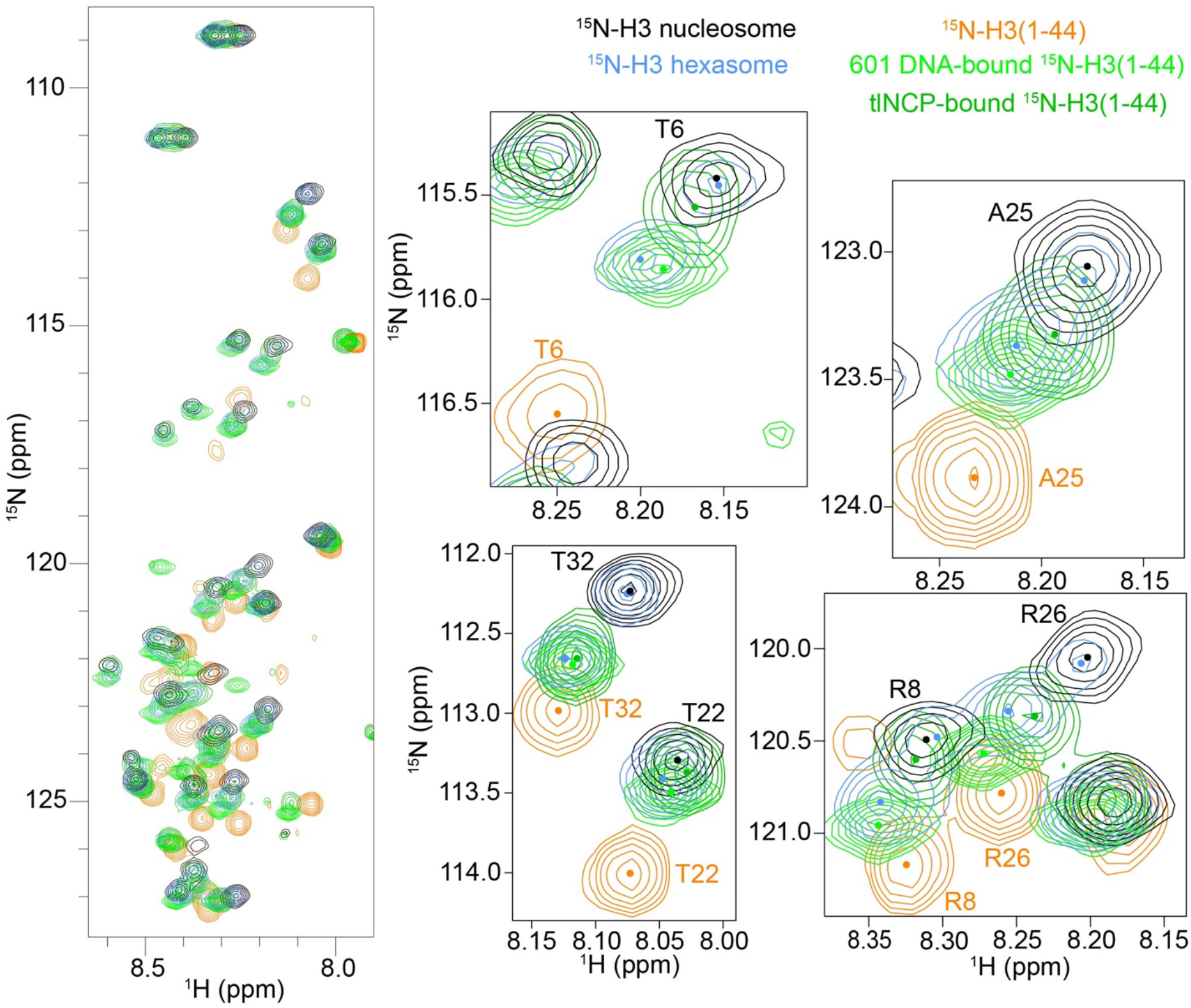
Same as main text Figure 3, except that spectra are additionally overlaid that were collected on ^15^N-H3(1-44) bound to either 601 DNA (lime green) or tailless NCP (green, from ^32^).

**Figure 4—Figure supplement 1.**
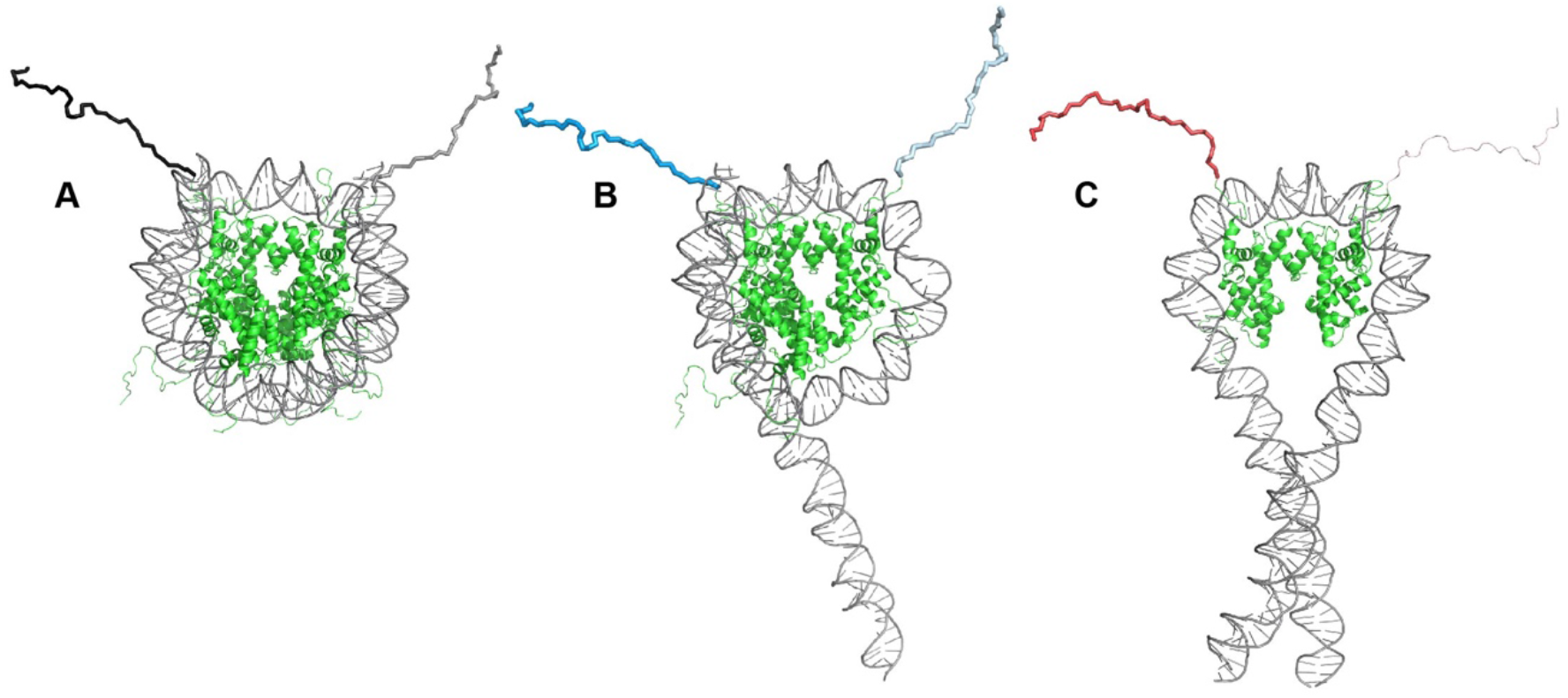
Starting states of nucleosomes and subnucleosomes for MD simulations. Initial states are shown for each of (A) nucleosome, (B) hexasome, and (C) tetrasome. The tail colors/shades of the H3 tails correspond to main text Figure 4.

**Figure 4—Figure supplement 2.**
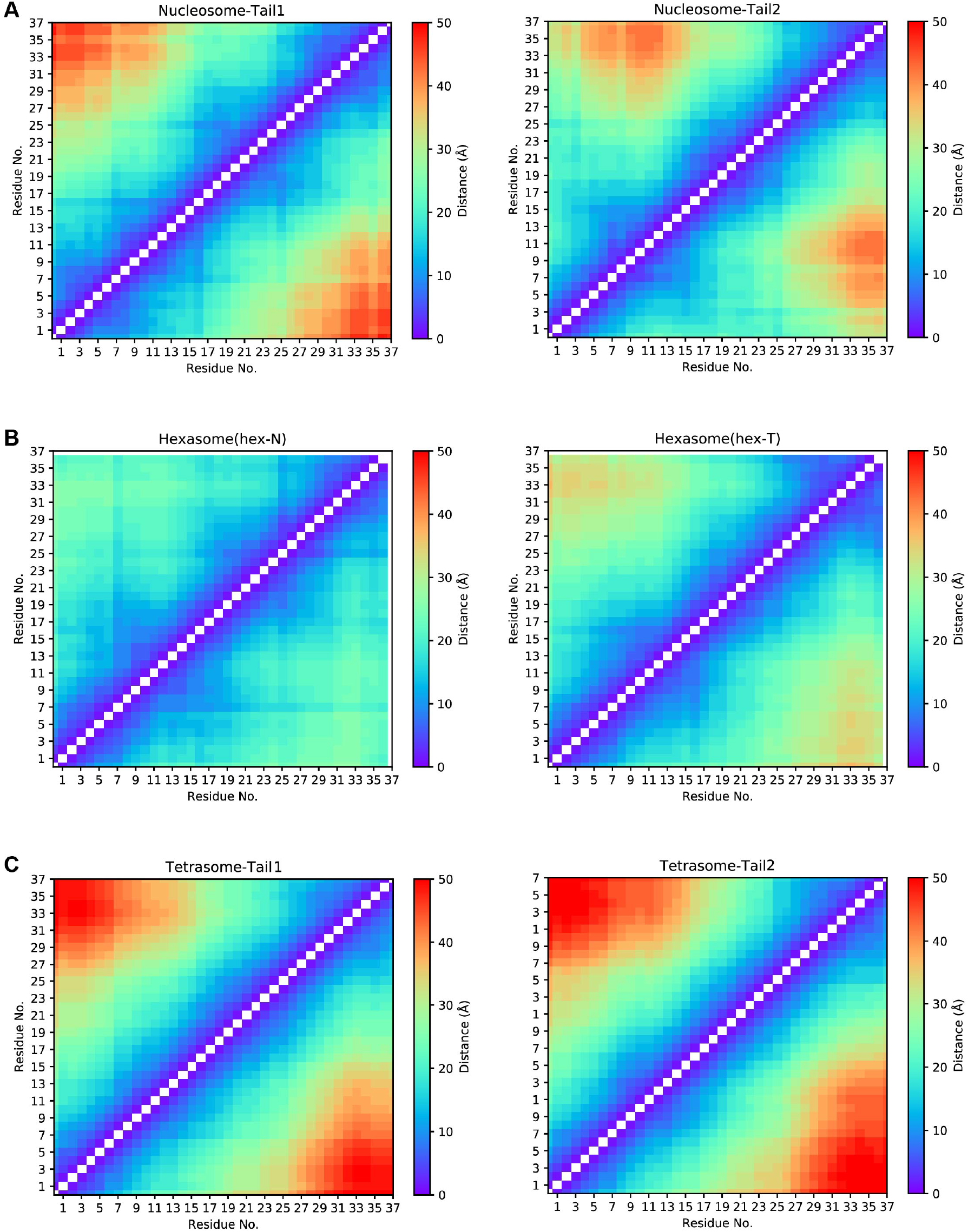
Calculated average intra-tail distances along the H3 tails from MD simulations. Plots are shown for each of (A) nucleosome, (B) hexasome, and (C) tetrasome. These plots report on compactness of the H3 tails.

**Figure 7—Figure supplement 1.**
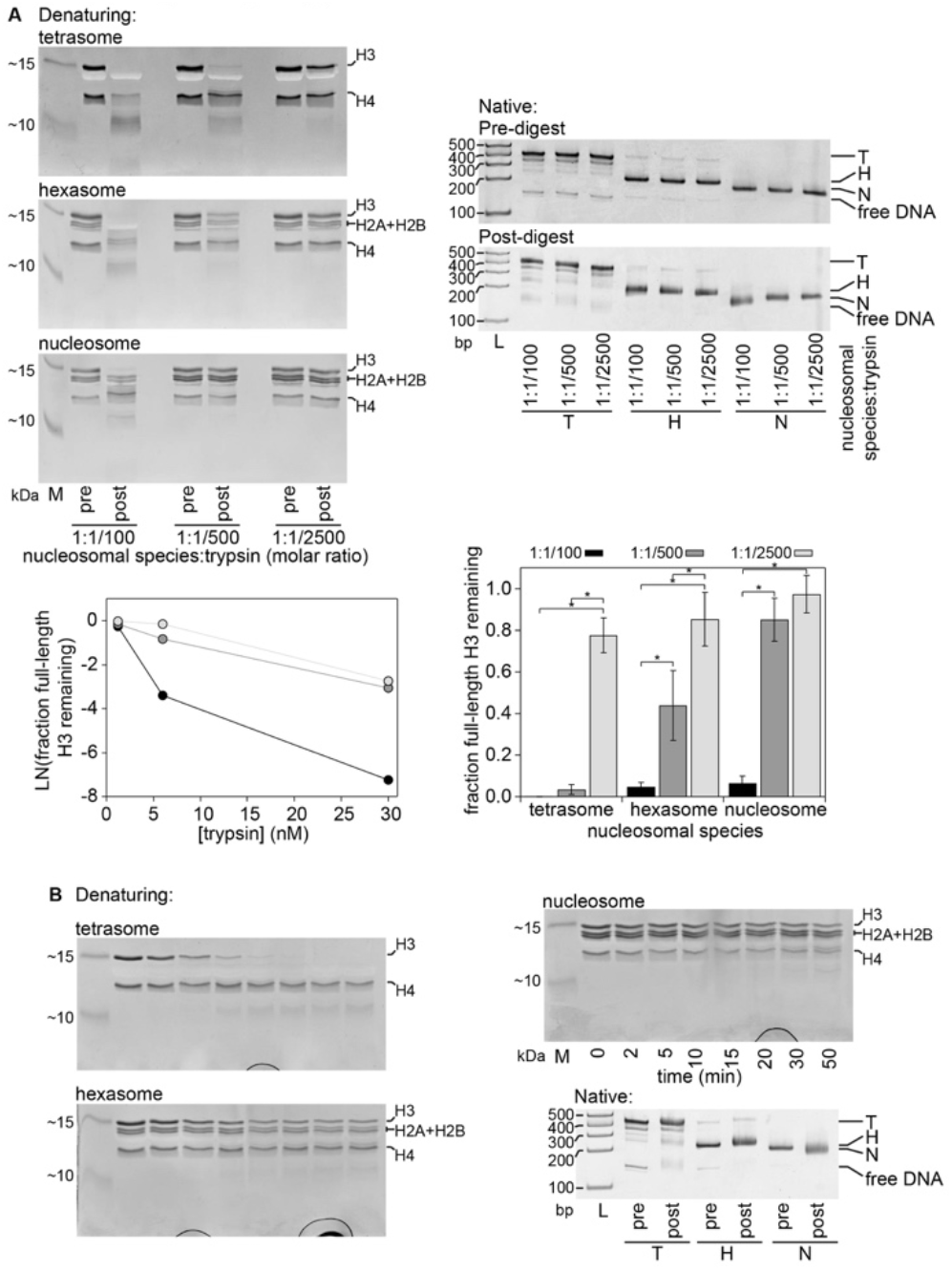
Trypsin digestion assays to probe tail accessibility. **A.**18% denaturing acrylamide gels (left) were used to assess the progression of trypsin proteolysis. Gel samples were taken before the addition of trypsin (t=0, “pre”) and 20 minutes after the addition of trypsin (t=20min, “post”) for three different molar ratios of nucleosomal species:trypsin and a fixed concentration of the given nucleosomal species (3μM). Quantification of the gels are shown in **Figure 7A**. 5% native acrylamide gels (right) confirm that the nucleosomal species remain intact during the assay. The amount of free DNA does not appear to increase significantly while the band for each nucleosomal species remains intact but appears to blur as a result of the tail cleavage. An alternative representation of **Figure 7A** (lower right) highlights differences between ratios of trypsin for a given nucleosomal species that are statistically significant as determined by a two-way ANOVA followed by a tukey post-hoc analysis (*, p<0.05). Plotting the natural log of the fraction of full length H3 remaining as a function of trypsin concentration (lower left) suggests a linear relationship between k_obs_ and enzyme concentration (see Materials and Methods for more details). **B.** 18% denaturing acrylamide gels (left) were used to assess the progression of trypsin proteolysis as a function of time at 3μM of a given nucleosomal species and a 1:1/500 molar ratio of trypsin. Gel samples were taken before the addition of trypsin (t=0) and at the indicated timepoints after the addition of trypsin. Quantification of the gels are shown in **Figure 7B**. 5% native acrylamide gels (lower right) confirm that the nucleosomal species remain intact during the assay. In the denaturing gels, the full-length position of each histone is labeled to the right of the gel. The denaturing gels were stained with Coomassie and include Spectra BR marker (M) for size reference while the native gels were visualized with ethidium bromide and include TrackIt 100bp DNA ladder (L) for size reference.

## Acknowledgements

This work in the Musselman group was funded by an NSF CAREER Award (1452411) and an NIH NIGMS R35 award (R35GM 128705). EAM was supported in part by an Arnold O. Beckman Postdoctoral Fellowship and by the Medical College of Wisconsin. This work in the Wereszczynski group was supported by an NSF CAREER Award (1552743) and an NIH NIGMS R35 award (R35GM119647). Work in the Poirier group is supported by the NIH (GM131626 and GM121966). The content is solely the responsibility of the authors and does not necessarily represent the official views of the National Institutes of Health. This work used the Extreme Science and Engineering Discovery Environment (XSEDE^81^), which is supported by National Science Foundation Grant No. ACI-1053575. We would like to thank the Carver College of Medicine and MCW (supported by NIH award number S10OD025000) NMR facilities. We would additionally like to thank the High Resolution Mass Spectrometry Facility (Office of the Vice-President for Research and Economic Development at the University of Iowa). Thanks also to Dr. John Egner for help with R analysis. Additional thanks to Drs. Karolin Luger for the gifts of the histone plasmids. Thanks to Dr. Sam Bowerman and Dr. Srinivas Ramachandran for helpful discussions.

## Competing Interests

The authors have no competing interests to disclose.

## Notes

### Competing Interest Statement

The authors have declared no competing interest.

